# Multi-stage resistance to *Zymoseptoria tritici* revealed by GWAS in an Australian bread wheat (*Triticum aestivum* L.) diversity panel

**DOI:** 10.1101/2022.06.29.498182

**Authors:** Nannan Yang, Ben Ovenden, Brad Baxter, Megan C. McDonald, Peter S. Solomon, Andrew Milgate

## Abstract

Septoria tritici blotch (STB) has been ranked the third most important wheat disease in the world, threatening a large area of wheat production. Although major genes play an important role in the protection against *Zymoseptoria tritici* infection, the lifespan of their resistance unfortunately is very short in modern agriculture systems. Combinations of quantitative resistance with minor effects, therefore, are believed to have prolonged and more durable resistance to *Z. tritici*. In this study new quantitative trait loci (QTLs) were identified that are responsible for seedling-stage resistance and adult-plant stage resistance (APR). More importantly was the characterisation of a previously unidentified QTL that can provide resistance during different stages of plant growth or multi-stage resistance (MSR). At the seedling stage, we discovered a new isolate-specific QTL, QSt.wai.1A.1. At the adult-plant stage, the new QTL QStb.wai.6A.2 provided stable and consistent APR in multiple sites and years, while the QTL QStb.wai.7A.2 was highlighted to have MSR. The stacking of multiple favourable MSR alleles was found to improve resistance to *Z. tritici* by up to 40%.

**Key message:** An Australian GWAS panel discovered three new QTLs associated with seedling-stage resistance, adult-plant stage resistance, and multi-stage resistance, respectively.

## Introduction

*Zymoseptoria tritici* (*Mycosphaerella graminicola* (Fuckel) J. Schrot, anamorph *Septoria tritici*, synonym) (Quaedvlieg et al. 2011), severely threatens wheat production in Australia, Europe, and North America. STB has been documented as the third most important disease threatening wheat production with average of 2.44% yield losses per year (Savary et al. 2019). Thirty to fifty percent yield loss is possible in regions that experience high humidity and mild temperatures during the growing season (Eyal 1987). The use of fungicides to control the spread of *Z. tritici* is becoming more challenging, due to the increasing levels of resistance to azole fungicides (McDonald et al. 2019; Milgate 2014) and recently strobilurin resistance observed in Australia (pers. comm F. Lopez-Ruiz). Resistance to multiple fungicide chemistries occurs in Europe and the US, including azole, strobilurin (Hagerty et al. 2017) and succinate dehydrogenase inhibitors (SDHI) (Blake et al. 2018; Dooley et al. 2016). To reduce the instance of fungicide resistance, fungicides need to be used as a part of an integrated disease management (IDM) system. An important IDM component for growers is having a robust level of host resistance to *Z. tritici* in cultivated wheat varieties.

Major resistance (R) genes are important sources of resistance that wheat breeders can use to protect against *Z. tritici*. To date, twenty-four *Z. tritici* major resistance genes have been reported with tightly linked molecular markers, including 12 isolate-specific genes and 12 non-isolate specific genes from wheat (Aouini 2018; Langlands-Perry et al. 2022; Tabib Ghaffary et al. 2018; Yang et al. 2018). However, this fungus has a plastic number of chromosomes, sexual and asexual reproduction systems, and the ability of long-distance migration, which increases the threat of the host resistance being overcome (McDonald and Mundt 2016; Rudd 2015). For instance, the major seedling resistance and adult-plant resistance (APR) genes, Stb4 (Adhikari et al. 2004), Stb6 (Brading et al. 2002), Stb2/WW/Stb11 (Dreisigacker et al. 2015; Liu et al. 2013; Raman et al. 2009), and Stb18 (Tabib Ghaffary et al. 2011) have been overcome in Australia (pers. comm A. Milgate). Thus, new sources of resistance are urgently required by breeders.

Quantitative resistance has been proposed as an alternative and potentially more durable solution to control STB. Different types of quantitative resistance have been identified in 89 genomic regions in wheat, of which, 27 were detected at the seedling stage, and 48 at the adult stage (Brown et al. 2015; Goudemand et al. 2013). New quantitative trait loci (QTLs) at the adult-plant stage have also been detected from nine GWAS studies (Alemu et al. 2021; Arraiano and Brown 2017; Kidane et al. 2017; Louriki et al. 2021; Mahboubi et al. 2022; Muqaddasi et al. 2019; Riaz et al. 2020; Vagndorf et al. 2017; Würschum et al. 2017; Yates et al. 2019). These include notable loci such as QStb.NS-2A associated with APR (Vagndorf et al. 2017), qtl-3 on the control of necrosis lesions (Yates et al. 2019), and QStb.teagasc-4A.1 associated with the STB resistance of flag leaves and flag-1 leaves (Riaz et al. 2020). However, there is a paucity of reports demonstrating the deployment of quantitative genes for *Z. tritici* resistance that can provide stable protection over a long period of time. On the other hand, STB levels can also be reduced by traits such as taller plant and late heading date or flowering time that contribute to disease escape, which limits the spread of fungal inoculum within crops (Arraiano et al. 2009; Brown 2015; Simón et al. 2004; Tavella 1978), although these traits may be unfavourable in breeding.

Unlike the seedling resistance observed in wheat leaf rust (*Puccinia triticina*), which often provides complete resistance at all growth stages (Ellis et al. 2014; Wellings et al. 2012). The translation of seedling resistance against *Z. tritici* infection to the adult plant is seldomly observed. Also, unlike the rust APR genes such as Lr34 and Lr46 (Lagudah et al. 2009; Sivasamy et al. 2014), which have provided durable resistance at the adult-plant stage, there is little evidence from the literature this would be the case for *Z. tritici*. Therefore, the combination of different seedling-resistance and APR has long been proposed as a sustainable way of prolonging the durability of disease resistance (Brown 2015; Niks et al. 2015; Rimbaud et al. 2021). Here we introduce the term “multi-stage resistance” (MSR) to describe those QTLs which are effective at more than one plant growth stage. This is defined as QTLs that reduce disease during both seedling stage and adult-plant growth stages. QTLs with MSR, which continue providing the resistance from early stages to later stages of the plant growth will be very desirable breeding targets. In this study, a collection of 273 bread wheat cultivars that represented both the gene pool of recent Australian cultivars and international sources of resistance were applied in a marker and trait genome-wide association analysis.

## Materials and methods

### Plant materials

Two hundred and seventy-three accessions were selected for inclusion into the AusSTB diversity panel. The panel is comprised of 163 cultivars and breeding lines from breeding programs across Australia, and a selection of *Z. tritici* resistance sources from around the world that are relevant to Australian *Z. tritici* resistance breeding. These include 13 synthetic hexaploid lines, 59 accessions from CIMMYT, 12 from North America, 10 from Europe, five from the Middle East, three each from New Zealand and Mexico, two each from Brazil and China and one from Russia. (Table S1). Accessions were sourced from the Australian Grains Genebank, Horsham Victoria Australia, and accession numbers are also provided in Table S1. Accessions were subjected to two generations of single seed decent to decrease genetic heterozygosity prior to phenotyping and DNA extraction.

### Experimental design

The package DiGGer (Coombes 2002) in R (R Core Team 2020) was used to create randomized complete block (spatial) designs for all experiments in the study. For the glasshouse experiments, the 273 AusSTB lines and six control cultivars ‘M1696’ (resistant, R) ‘Teal’ (R), ‘Milan’ (moderately resistant/moderately susceptible, MR/MS), ‘Millewa’ (MR/MS), ‘Egret’ (susceptible, S), and ‘Summit’(S) were replicated three times in a 30 row by 30 column array in each experiment. For the field experiments, the 273 AusSTB lines were replicated three times and the balance of entries in each experiment were made up of the susceptible control cultivar ‘WW425’. The spatial designs for field experiments were 28 row-by-30 column arrays at Wagga Wagga New South Wales (NSW) and 12 rows by 69 columns arrays in Hamilton Victoria (VIC).

### Glasshouse screening for *Z. tritici* resistance

Three Australian *Z. tritici* isolates were used in this study. WAI332 was collected from NSW in 1979, WAI251 from VIC in 2012 and WAI161 from Tasmania (TAS) in 2011. Inoculation procedure for the isolates and experimental details for phenotyping *Z. tritici* glasshouse infections are described in Yang et al. (2018). Each isolate was screened on all 273 cultivars in six independent glasshouse screening experiments.

Symptoms of *Z. tritici* were assessed between 21 and 28 days after inoculation. A seedling infection score (STB_S) was scored based on the visually estimated percentage of necrotic lesions containing pycnidia on the infected leaves, based on the methods in Zwart et al. (2010). During the assessment, the percentage of leaf area with necrosis (Nec, 0-100%) on the infected leaf and pycnidia density on the necrotic leaf area (Pyc, 0-100%) were also recorded.

### Field experiments

The AusSTB panel was evaluated in four different environments (two locations × two years). Field experiments were conducted at Wagga Wagga Agricultural Institute at Wagga Wagga, NSW, Australia (WGA, −35.04419222, 147.3167896) in 2015 and 2016, and the Department of Economic Development, Jobs, Transport and Resources Hamilton Centre at Hamilton, Victoria, Australia (HLT, −37.828768, 142.082319) in 2015 and 2016. For the field experiments at Wagga Wagga, natural infection of *Z. tritici* was supplemented with an inoculation of stubble debris from a local crop with known high level of *Z. tritici* infection. Field experiments at Hamilton relied on natural infections.

Disease severity in the field experiments was visually scored according to Saari and Prescott’s severity scale for assessing wheat foliar diseases (Saari and Prescott 1975). Namely, STB_A (1-9) was used to record the observations. STB_A (1-9) is used to record *Z. tritici* disease intensity considering the plant growth stage, while STB (1-00%) is used to reflect the disease severity by recording the proportion of plant units with diseased leaves.

Two phenotypic scores were collected at Wagga Wagga in 2015, approximately four weeks apart in mid-September (Cycle 1, C1) and mid-October (Cycle 2, C2). One score collected at all other field experiments in mid-October. Additionally, relative maturity was scored using the Zadoks growth scale (Zadoks et al. 1974) at the same time as disease scores were collected. Plant height (HT) measurements of each entry was collected in 2016 and 2017 on December at physiological maturity.

### DNA extraction and genotyping

Leaf tissue was harvested from 14-day old seedlings and used for DNA extraction. DNA extraction and genotyping service were conducted by DArT Pty Ltd, Canberra, ACT. Genetic positions of all the markers were assigned according to the custom Chinese Spring Consensus Wheat map v4.0 provided by DArT (pers. comm Dr Andrzej Kilian).

The 273 DNA samples from each line in the population were assayed with two technical replications to derive reproducibility scores. At the first-stage quality control, the reproducibility rate was 0.95 for SNPs and 0.99 for silicoDArTs, and the call rate was 0.85 for SNPs and 0.95 for silicoDArTs. Details of the experimental procedure for generating silicoDArTs are described by Courtois et al. (2013) and Li et al. (2015). At the second stage of quality control, duplicated markers, markers with a Minor Allele Frequency (MAF) < 0.05, and markers not assigned to the chromosome map were excluded. The final marker sets for the association study comprised of 11,200 SNPs and 29,346 silicoDArTs (Table 1).

**Table 1.** Summary of the average of Polymorphism Information Content (PIC), the average of Minor Allele Frequency (MAF), and the number of SNP and silicoDArT markers on each chromosome

| Chr | Genetic Length (cM) | SNP |  |  | silicoDArT |  |  |
| --- | --- | --- | --- | --- | --- | --- | --- |
|  |  | Avg. PIC | Avg. MAF | No. of markers | Avg. PIC | Avg. MAF | No. of markers |
| 1A | 255.6 | 0.28 | 0.26 | 617 | 0.31 | 0.26 | 1,113 |
| 2A | 138.6 | 0.24 | 0.25 | 752 | 0.3 | 0.26 | 1,836 |
| 3A | 154.2 | 0.26 | 0.25 | 674 | 0.31 | 0.26 | 1,208 |
| 4A | 135.2 | 0.25 | 0.25 | 540 | 0.31 | 0.27 | 1,612 |
| 5A | 160 | 0.26 | 0.26 | 568 | 0.31 | 0.28 | 847 |
| 6A | 105 | 0.22 | 0.29 | 526 | 0.26 | 0.26 | 1,344 |
| 7A | 160.2 | 0.23 | 0.25 | 658 | 0.3 | 0.27 | 1,739 |
| A | 1,108.8 | 0.25 | 0.26 | 4,335 | 0.3 | 0.27 | 9,699 |
| 1B | 286.6 | 0.21 | 0.24 | 822 | 0.25 | 0.21 | 2,740 |
| 2B | 110 | 0.22 | 0.22 | 1,468 | 0.26 | 0.21 | 4,377 |
| 3B | 161.1 | 0.25 | 0.27 | 879 | 0.31 | 0.26 | 2,351 |
| 4B | 86.3 | 0.25 | 0.27 | 314 | 0.31 | 0.28 | 591 |
| 5B | 153.7 | 0.28 | 0.29 | 1,041 | 0.34 | 0.29 | 2,134 |
| 6B | 87.9 | 0.24 | 0.26 | 612 | 0.29 | 0.27 | 1,767 |
| 7B | 142 | 0.24 | 0.25 | 485 | 0.3 | 0.26 | 1,784 |
| B | 1,027.6 | 0.24 | 0.26 | 5,621 | 0.29 | 0.25 | 15,744 |
| 1D | 139.5 | 0.21 | 0.24 | 244 | 0.3 | 0.29 | 548 |
| 2D | 166.2 | 0.21 | 0.21 | 400 | 0.3 | 0.24 | 1,351 |
| 3D | 156.1 | 0.13 | 0.21 | 132 | 0.28 | 0.27 | 665 |
| 4D | 97.4 | 0.15 | 0.22 | 52 | 0.27 | 0.25 | 152 |
| 5D | 154.2 | 0.17 | 0.28 | 107 | 0.29 | 0.26 | 311 |
| 6D | 112.5 | 0.18 | 0.27 | 131 | 0.28 | 0.29 | 379 |
| 7D | 190.7 | 0.14 | 0.25 | 178 | 0.24 | 0.26 | 497 |
| D | 1,016.6 | 0.17 | 0.24 | 1,244 | 0.28 | 0.27 | 3,903 |
| **Total** | **3,153** | **0.22** | **0.25** | **11,200** | **0.29** | **0.26** | **29,346** |

### Linkage disequilibrium and population structure analysis

The R package LDheatmap (Shin et al. 2006), was used to obtain the linkage disequilibrium (LD) squared allelic correlation (*r*^2^) estimates for all pairwise comparisons between markers on each chromosome for each marker set separately. To quantify the pattern of LD decay, syntenic pairwise LD *r*^2^ estimates were plotted against the corresponding pairwise genetic distances for each of the A, B and D genomes and for the overall wheat genome. A second degree locally weighted polynomial regression (LOESS) curve was fitted to each scatter plot (Cleveland and Devlin 1988) following the approach of Maccaferri et al. (2015). The intersection of the LOESS curve and an *r*^2^ threshold of 0.20 for marker pairs was taken as an estimate of the extent of LD decay within each genome for each marker set and was used to define the confidence intervals of QTL detected in this study (Fig. S3).

Population structure of the AusSTB panel was analysed using the software package STRUCTURE version 2.3.4 (Pritchard et al. 2000), using both SNP and silicoDArT marker sets. An admixture model with 10 predefined subpopulations replicated 10 times was run with 10,000 iterations of burn-in followed by 10,000 recorded Markov-Chain iterations for each marker set. Output from STRUCTURE was analysed in the R package Pophelper (Francis 2017) to determine the optimal number of subpopulations using the Evanno method (Evanno et al. 2005). STRUCTURE was then re-run with the optimum number of subpopulations (seven) to generate population membership coefficient matrices (***Q***) as well as the corresponding population membership coefficients, were obtained for each marker set (Fig. 1).

**Figure 1.**
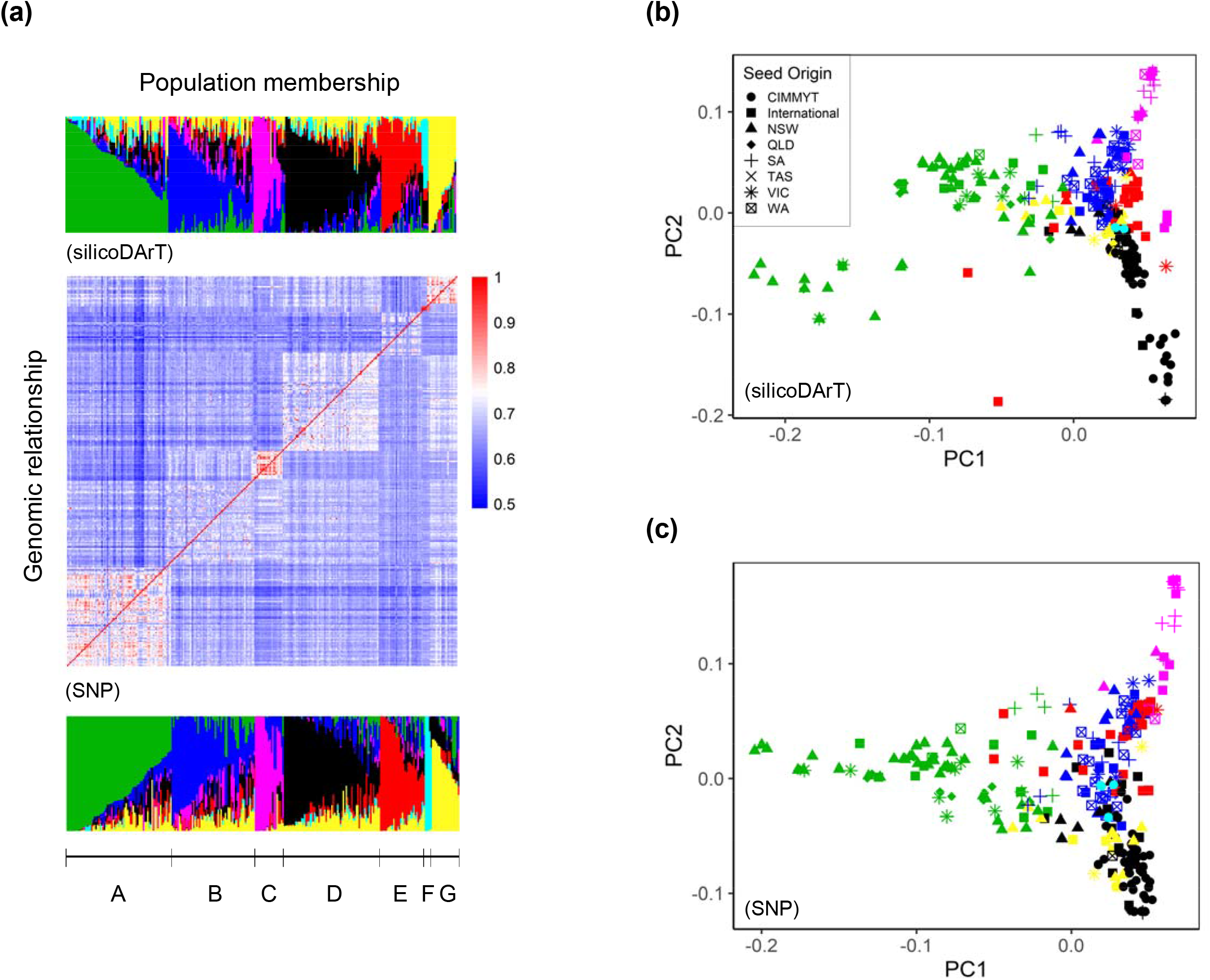
Population structure analysis of the AusSTB panel. **(a)** Genomic relationship matrix (***K***) and population membership coefficient matrices (***Q***) showing the seven hypothetical subpopulations derived from the STRUCTURE analysis. **(b)** Principal components analysis of the AusSTB panel using the silicoDArT markers. **(c)** Principal components analysis of the AusSTB panel using the SNPs. The seven sub-populations are displayed in Green (A), Blue (B), Pink (C), Black (D), Red (E), Cyan (F), and Yellow (G). Seed origins of different accessions are CIMMYT (•), International (◼), NSW (▲), QLD (♦), SA (+), TAS (×), VIC (*), and WA (⊠), respectively.

### Phenotypic data analysis

A multiplicative mixed linear model was used to analyse phenotype data for each trait at each experiment following the approach of Gilmour et al. (1997), using the R software package ASReml-R version 3 (Butler et al. 2018), in the R statistical software environment (R Core Team 2020). The linear mixed model is given by

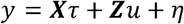

where *y* is the (*n* × 1) data vector of the response variable; *τ* is a (*t* × 1) vector of fixed effects (including genetic line effects and the intercept) with associated design matrix ***X***. The term *u* is a random component with associated design matrix ***Z*** and contains the experimental blocking structures (replicate, range and row) used to capture extraneous variation. Random effects were maintained in the model if they were significant according to log likelihood ratio tests relative to the full model (Stram and Lee 1994). The residual error is *η* was assumed to have distribution *η* ∼ *N*(0, *σ*^2^***R***) where *σ*^2^ is the residual variance for the experiment and ***R*** is a matrix that contains a parameterization for a separable autoregressive AR1 ⨂ AR1 process to model potential spatial correlation of the observations.

A total of 31 models were constructed for the traits collected from six experiments in GH and four experiments in the field (Table S2). Best Unbiased Linear Estimates (BLUEs) were obtained from each model for subsequent use in the association analyses.

### Genome-wide association analysis

Association analyses using the phenotype BLUEs described above were performed using the R software package Genome Association and Prediction Integrated Tool (GAPIT) version 3 (Tang et al. 2016). Missing markers in the two marker sets (consisting of 11,200 SNPs and 29,346 silicoDArTs) were imputed with the major allele at each locus using the imputation function in GAPIT. A separate scaled identity by descent relationship matrix (***K***) after VanRaden (2008) was calculated for each marker set. Separate association analyses for each trait in each experiment and for the two different marker sets were performed using the compressed mixed linear model approach (Zhang et al. 2010), implemented in GAPIT as follows:

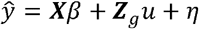

where *ŷ* is the vector of BLUEs for one trait measured in one experiment, *β* is a vector of fixed effects for the corresponding design matrix (***X***), including the molecular marker, population assignments from the STRUCTURE analysis (***Q***) and the intercept. The vector of overall genetic line effects u (with associated design matrix ***Z***_*g*_) is modelled as 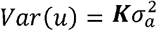, where ***K*** is the relationship matrix and 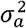 is the estimated additive genetic variance. *η* is the vector of random residuals.

Genetic regions that had marker-trait associations at False discovery rate (FDR) adjusted *p*-value less than 0.3 and were also detected by at least two GWAS were considered QTLs (Maccaferri et al. 2015; Nyine et al. 2019; Ovenden et al. 2017), as repeated detection provides more support for biological association. The confidence interval for QTL is calculated from the genome-wide LD threshold determined above: 1-4 cM for the A and B genomes and 4-6 cM for the D genome. The marker with the lowest *p*-value at each QTL was considered the representative marker for the QTL. In addition, to compare the differences among different groups of stacking alleles, Wilcoxon Rank Sum tests was used to generate the *p* values.

### Bioinformatic analysis

The 13 QTLs identified in this study were compared to over 100 QTLs from six GWAS studies, studies using segregating populations and 24 named *Z. tritici* genes (Aouini 2018; Langlands-Perry et al. 2022; Tabib Ghaffary et al. 2018; Yang et al. 2018). Firstly, genomic DNA sequences of the 13 QTLs based on the confidence intervals were extracted from the IWGSC RefSeq v1.0 (https://urgi.versailles.inra.fr/). Secondly, the DNA sequences of Simple Sequence Repeats (SSR), silicoDArT and SNP markers tightly linked to reported QTLs were search against the physical QTL regions using blastn. Only those QTLs that overlapped or were detected in our QTL regions were considered as co-localization. All the Coding Sequences (CDS) were then extracted from the 14 QTL regions, and then were BLAST against 314 representative annotated R genes from wheat, maize, rice, and Arabidopsis (Kourelis and van der Hoorn 2018) to identify candidate R genes in the QTL reported in this study. The KASP marker wMAS000033, provided by Integrated Breeding Platform (IBP, https://www.integratedbreeding.net/), was used to track the allele frequency of the gene *Vrn-1A* in the AusSTB panel.

## Results

### Genotypic data and LD estimation of the AusSTB panel

The consensus map contained all 21 bread wheat chromosomes, covering 3,153 cM, with a total number of 11,200 SNPs and 29,346 silicoDArTs (Table 1). Overall, silicoDArTs were 2-5 times more frequently detected than SNPs on all the 21 chromosomes. Although SNPs and silicoDArTs gave very similar patterns on each of the chromosome, the distribution of silicoDArTs had 9 less gaps than SNPs (Fig. S1). The average Minor Allele Frequency (MAF) was similar between SNPs (0.25) and silicoDArTs (0.26) on the A, B, and D genomes. The average Polymorphism Information Content (PIC) differed slightly between SNPs (0.24) and silicoDArTs (0.25) on A and B genomes, whereas the average PIC of SNPs (0.17) on D genome was 58% less than that of silicoDArTs (0.27).

Little difference in LD patterns was observed between SNPs and silicoDArTs on the 21 chromosomes, except for a few LD blocks on chromosome 1B, 1D, and 3D (Fig. S2). An overall average of genetic distance of LD decay was 3 cM for SNPs and 0.68 cM for silicoDArTs at *r*^*2*^ = 0.20 (Fig. S3c). When estimating LD decay of SNP and silicoDArT individually, SNP LD decay at *r*^*2*^ = 0.20 was 1.2 cM for the A genome, 4.4 cM for the B genome, and 8.5 cM for the D genome (Fig. S3a). Smaller LD blocks were captured by silicoDArTs than from SNPs in most genomic regions. LD decay values of silicoDArT, were 1.0 cM for the A genome, 0.7 cM for the B genome, and 2.3 cM for the D genome (Fig. S3b).

### AusSTB panel composition and genetic structure

The two sets of markers were used to calculate the genomic relationships matrix (***K***) and the Structure matrices (***Q***). This analysis indicated there were seven subpopulations amongst the 273 accessions. (Fig. 1a). The alignment of members in each subpopulation was not stable between SNPs and silicoDArTs from k = 2 to k = 6 (data not shown), until k = 7 where the discrepancy minimized, and the ***K*** matrix matched with the ***Q*** matrix (Fig. 1a). This population structure was strongly correlated with the seed origin based on the principal component analysis (PCA) (Fig. 1b and 1c). Specifically, 35 out of 73 members in Sub-population A originated from NSW, 19 out of 58 members in Sub-population B from Western Australian (WA), 8 out of 20 members in Sub-population C from South Australian (SA). The 31 members in the Sub-population E, originated from multiple places across the world, was clustered by their growth habits as winter or spring-winter (Table S1).

### Phenotypic data analysis of the AusSTB panel

The response of the 273 accessions to *Z. tritici* infection at the seedling stage were tested against three different *Z. tritici* isolates, which are representative pathotypes for south eastern Australia (WW332, WAI251 and WAI161). Normal frequency distributions were observed in this wheat panel for phenotypic traits Necrosis and STB_S, whereas the frequency distribution of Pycnidia phenotypes was skewed towards zero (Fig. 2a). The number of isolate-specific resistant accessions (STB_S scores, 1-2) varied from 23 (WAI161), 17 (WAI251), to 31 (WAI332). Fourteen wheat accessions (STB_S scores, 1-2) were resistant to three *Z. tritici* isolates in six independent experiments, and five of the most susceptible accessions (STB scores, 3.5-5) were also identified (data not shown).

**Figure 2.**
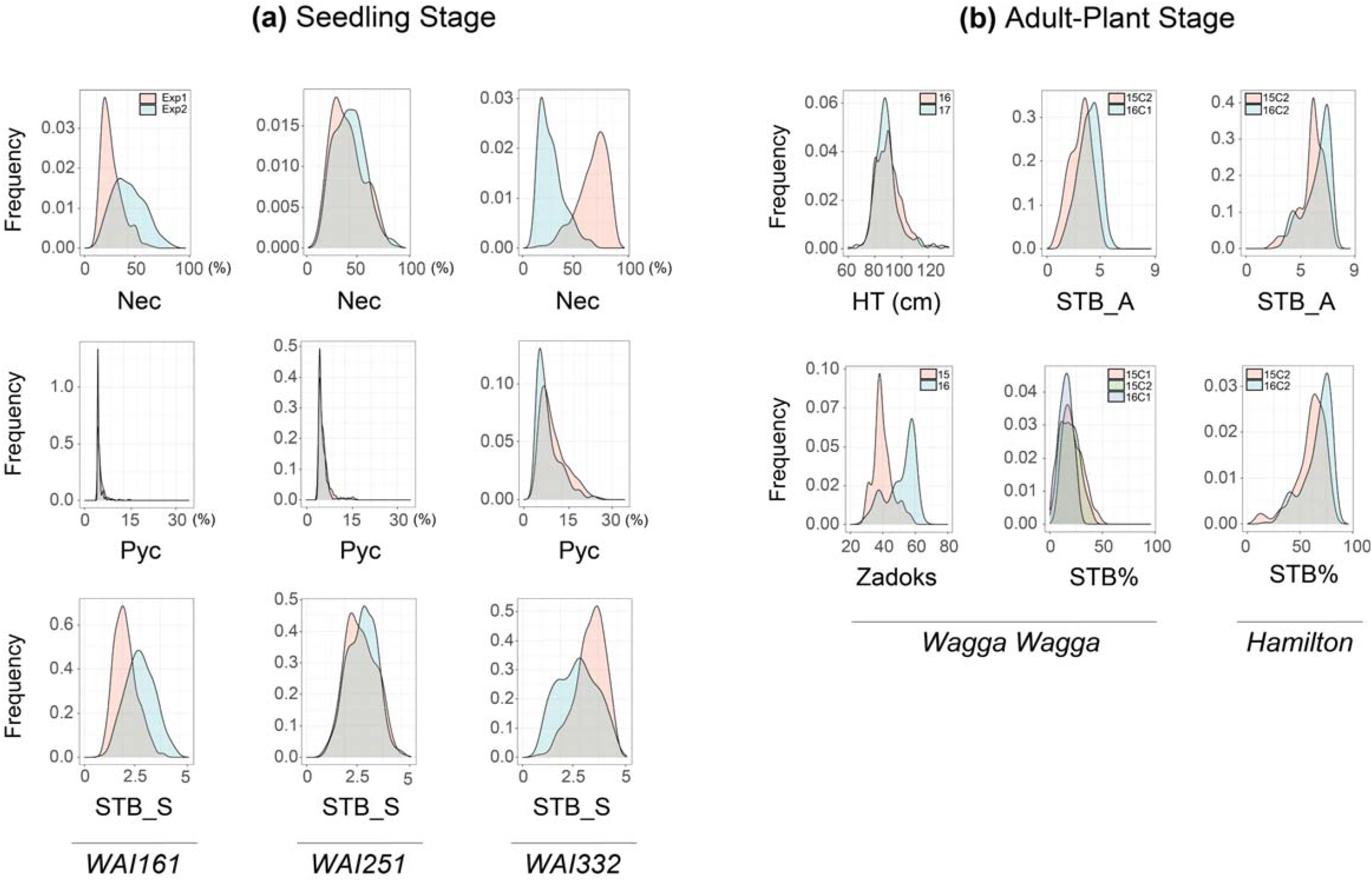
Frequency distributions of 31 BLUEs from the AusSTB panel. **(a)** Traits at the seedling stage include the percentage of necrotic leaf area (Nec) on the infected leaves, the pycnidia density (Pyc) in the necrotic leaf area, and the STB_S Scale 1 to 5 using the three STB isolates WAI161, WAI251, and WAI332. **(b)** Traits at the adult-plant stage include Plant Height (HT), Relative maturity (Zadoks scale), STB_A scale 1-9, and the percentage of STB infected leaf area on the whole plant (STB%). Traits HT and Zadoks were only measured at the site of Wagga Wagga.

The response of the 273 accessions to *Z. tritici* infection at the adult plant stage was evaluated in four environments (two locations x two years) under natural *Z. tritici* infection. High correlations were observed between four BLUEs of STB_A and five BLUEs of STB%, with Pearson correlation coefficient values ranging from *r* = 0.94 to 0.99 (Table S2). This suggests that the two scoring methods captured similar progress of *Z. tritici* infection on plants. A normal frequency distribution was observed on BLUEs of STB_A trait from Wagga Wagga, whereas the distribution shifted towards more susceptibility for the BLUEs of STB_A trait from Hamilton (Fig. 2b). Based on the adult-plant assessments in the AusSTB panel, none of the accessions displayed high levels of resistance (R), only five accessions were categorized as moderately resistant (MR). Approximately 20% of the accessions were categorized as being MSS, while the remaining accessions were categorized as susceptible (S) or susceptible/very susceptible (SVS, data not shown).

### Association analysis for *Z. tritici* resistance

Thirteen QTLs were detected at the seedling stage and at the adult-plant stage (Fig. 3, Table S3). These included six QTLs responsible for the *Z. tritici* isolate-specific resistance at seedling stage, which accounted for 3.8-6.9% phenotypic variance. Two QTLs were identified as non-isolate specific resistance as they were detected traits from more than two *Z. tritici* isolates (3.6-6.9% variance), one QTL at adult plant stage (3.2-4% variance), and four QTLs with multi-stage resistance (MSR, 3.1-6.7% variance).

**Figure 3.**
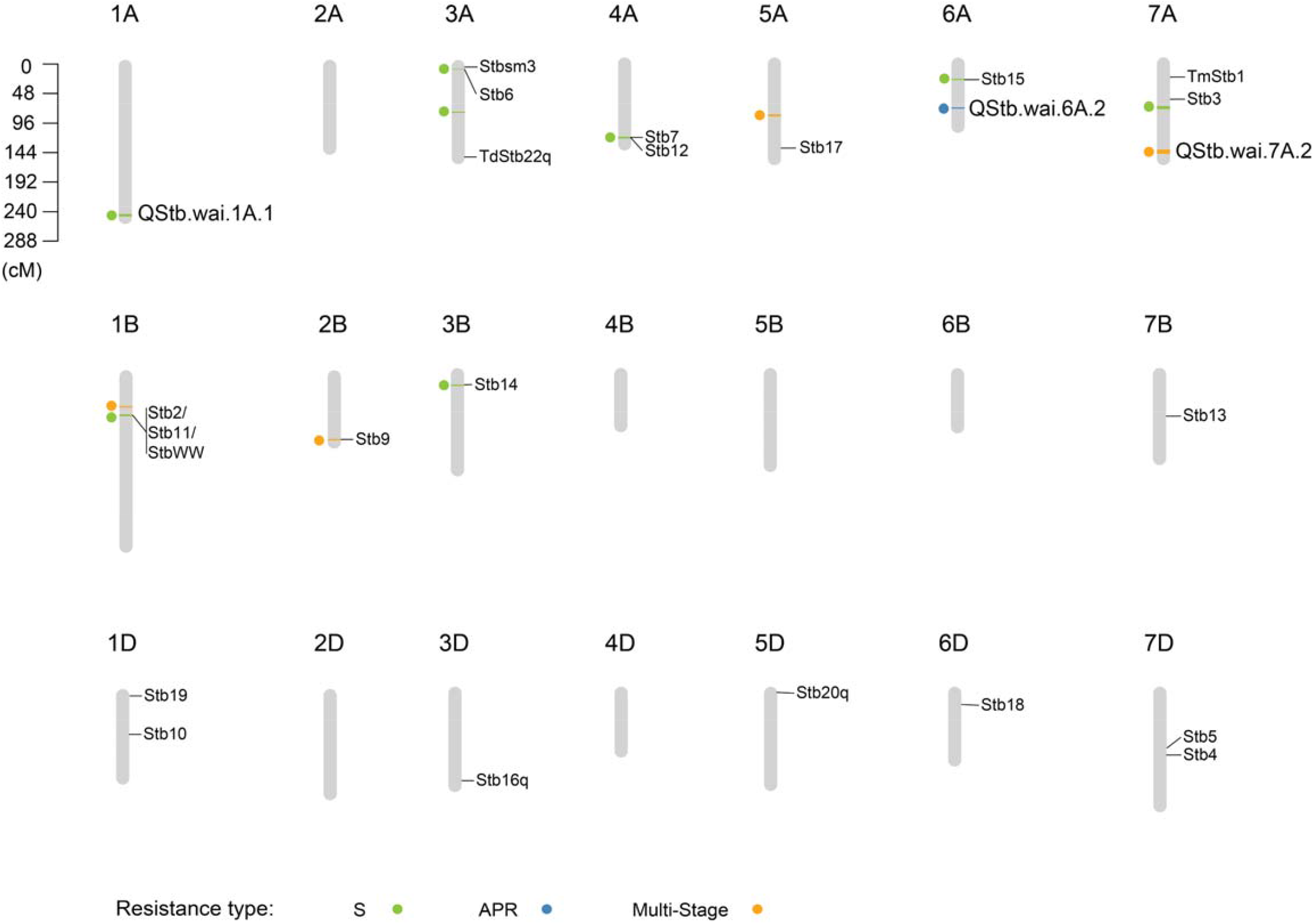
Genetic positions of detected QTLs associated with the *Zymoseptoria* resistance at the seedling stage and adult-plant stage. The 13 associations are shown as a solid circle on the left of each chromosome. A bar (linkage disequilibrium confidence interval) is shown in light green for the seedling stage resistance, blue for the adult-plant stage resistance, and orange for the multiple-stage resistance. Names and positions of the previously published *Zymoseptoria* major genes are also shown on the CS consensus genetic linkage map.

Three new QTLs were discovered in this study (Fig. 2 and Table 3), these are QStb.wai.1A.1 associated with *Z. tritici* resistance against the isolate WAI251, QStb.wai.6A.2 associated with APR and QStb.wai.7A.2 associated with MSR.

### *Z. tritici* resistance associated with HT and Zadoks traits

Plant height (HT) and plant growth stages (Zadoks) are two important phenological traits known to have various effects on the control of *Z. tritici* resistance. Slight to moderate negative correlations (*r* = −0.12 to −0.41, Table S2) were observed between HT and the 18 STB_A and STB% related BLUEs from the field data, suggesting shorter plants tended to have higher susceptibility. In contrast, Zadoks growth scale had strong positive correlations (*r* = 0.3-0.83, Table S2) with the 18 STB_A and STB% traits related BLUEs from the field data. The average of STB_A score in the Subpopulation E, which contained the most of the low-Zadoks scale and/or winter-type (slow-growing) plant accessions, was 15% lower than the other subpopulations (data not shown).

### QTLs associated with *Z. tritici* resistance at the seedling stage

Six QTLs were associated with the *Z. tritici* isolate-specific resistance, one with WAI161, one with WAI251, and four with WAI332 (Table 2). QStb.wai.6A.1 (resistant allele, C) was associated with the WAI161-specific resistance, with a resistant variant frequency of 0.83. Blast searches with our markers from the Chinese Spring reference genome indicated that QStb.wai.6A.1 co-located with the major gene Stb15 (Fig. 3). The new putative QTL, QStb.wai.1A.1 (resistant allele, **+**) was only detected from phenotypes recorded using inoculation with the isolate WAI251 (Fig. 3) and accounted for over 5% phenotypic variance. There are four WAI332-specific associated QTLs: QStb.wai.1B.2 (resistant allele, C) co-located with the major gene locus STB2/STB11/STBWW, QStb.wai.3A.1 (resistant allele, -) with Stb6, and QStb.wai.3B.1 (resistant allele, C) with Stb14 (Table 2, Fig. 3).Interestingly, QStb.wai.4A.1 (resistant allele, -), which resided in the Stb7/12 locus, was associated with traits HT and Zadoks, however there was insufficient data to further improve the resolution in this QTL region.

**Table 2.**
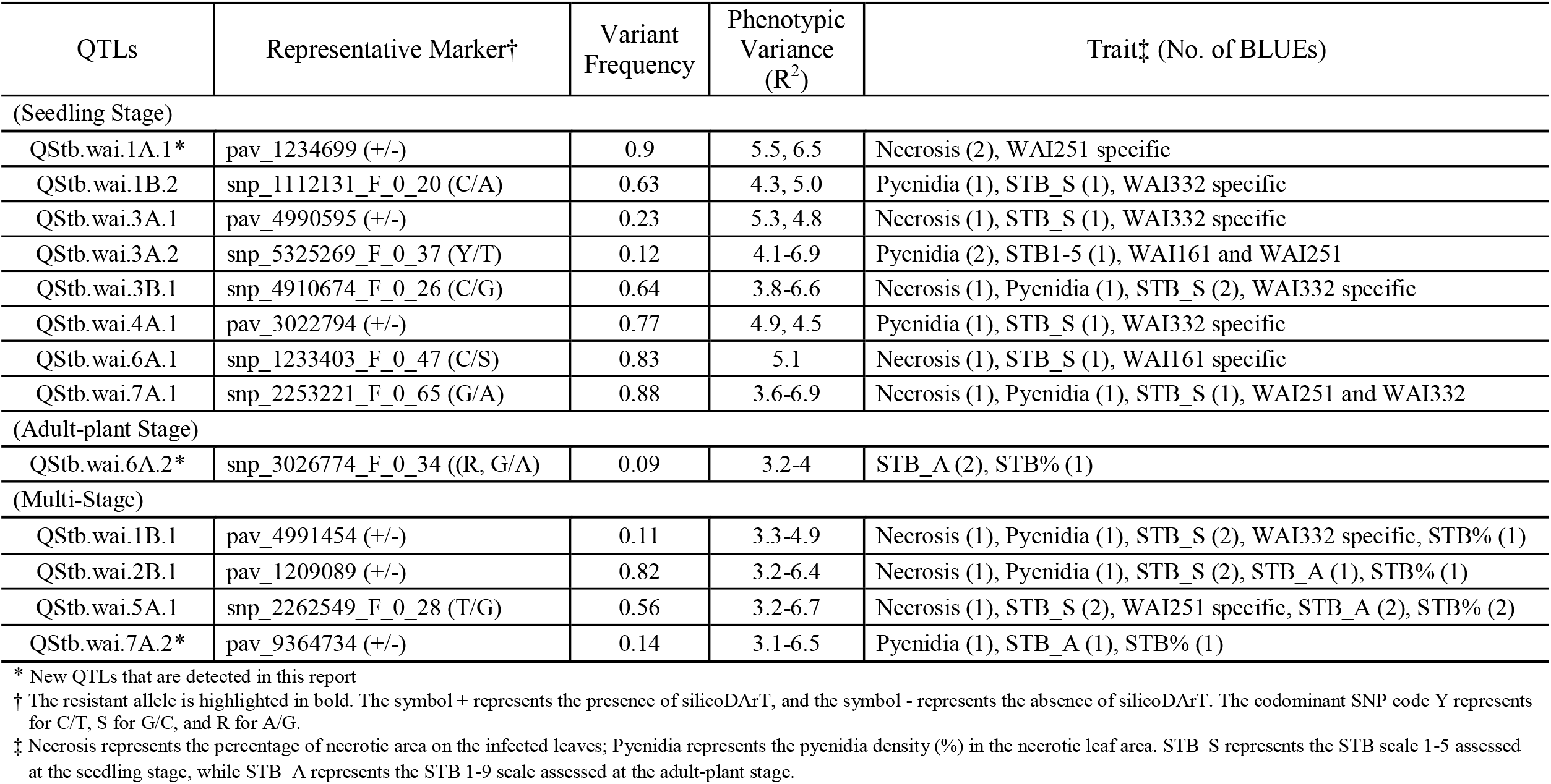
Summary of 13 QTLs detected by five traits associated with *Zymoseptoria tritici* resistance.

Two QTLs were detected as non-isolate specific resistance at the seedling stage (Table 2). The resistant QTL, QStb.wai.3A.2 (resistant allele, Y), was associated with pycnidia density, with a variant frequency of 0.12 (Table 2). QStb.wai.3A.2 was detected by phenotypes recorded using the isolates WAI161 and WAI251, with phenotypic variance ranging from 4.1-7.6%. The QTL, QStb.wai.7A.1 (resistant allele, G), also associated with three traits of WAI251 and WAI332, accounting for 3.6-6.9% phenotypic variance.

### QTLs associated with APR and MSR

Five QTLs associated with APR and/or MSR, having variant frequencies from 0.09 to 0.82 were identified (Table 2). These QTLs gave 3.1-6.7% phenotypic contributions to APR (Fig. 2).

The QTL, QStb.wai.6A.2, (resistant allele, R or G) is a new QTL, was confined in the region of 73-76 cM, associated with APR in 2015 and 2016 at both Wagga Wagga and Hamilton (Table 2). A blast search found that the tightly linked maker snp_3026774_F_0_34 peaked at the site of 454 Mb according to the Chinese Spring reference genome (S Table 3).

Four QTLs were categorized as MSR associated with multiple traits at the seedling stage and adult-plant stage (Table 2). The resistant QTL, QStb.wai.2B.1 (resistant allele, **+**), detected by non-specific isolate resistance at the seedling stage and the adult-plant stage, was found to collocate with the major gene Stb9 (Fig. 3). The second MSR QTL, QStb.wai.1B.1 (resistant allele, +) is close to QStb.wai.1B.2 associated with the resistance at the seedling stage, but our evidence suggests these are two separate QTLs. The genomic region of QStb.1B.1 spanned from 0 to 20 megabases (Mb), while QStb.wai.1B.2 was in the genomic region of 40 to 100 Mb (Table S3). Thirdly, QStb.wai.5A.1 (resistant allele, T) was highly associated with the Zadoks trait and APR (Table 2 and Fig. S4). Interestingly, the tagged SNP snp_2262549_F_0_28 of QStb.wai.5A.1 was also detected by two WAI251-seedling related traits, Necrosis and STB_S (Table 2). The fourth QTL, QStb.7A.2 (resistant allele, -) was defined in a genetic span of 4 cM on the distal region of 7AL associated with MSR (Fig. 4). The impact of MSR allele-stacking showed that combinations of QTL alleles with minor effects increased the overall resistance level in phenotypes recorded in this study. Combinations of three MSR alleles (**++T**+ and -**+T-**) showed superior performance, increasing the resistance by 10-30% at Hamilton and by 14-37% at Wagga Wagga (Fig. 5). Interestingly, the stacking of MSR alleles **++T**+ performed better (10% more resistance) than the stacking of -**+T-** at Hamilton, whereas the stacking of MSR alleles **++T**+ gave ∼5% less resistance than -**+T-** at Wagga Wagga. However, no significant differences (*p* values = 0.3) were observed between the combination **++T**+ of and the combination of -**+T-** at Hamilton and Wagga Wagga. Unfortunately, no accessions in the AusSTB panel had the combination of all four favourable MSR alleles together.

**Figure 4.**
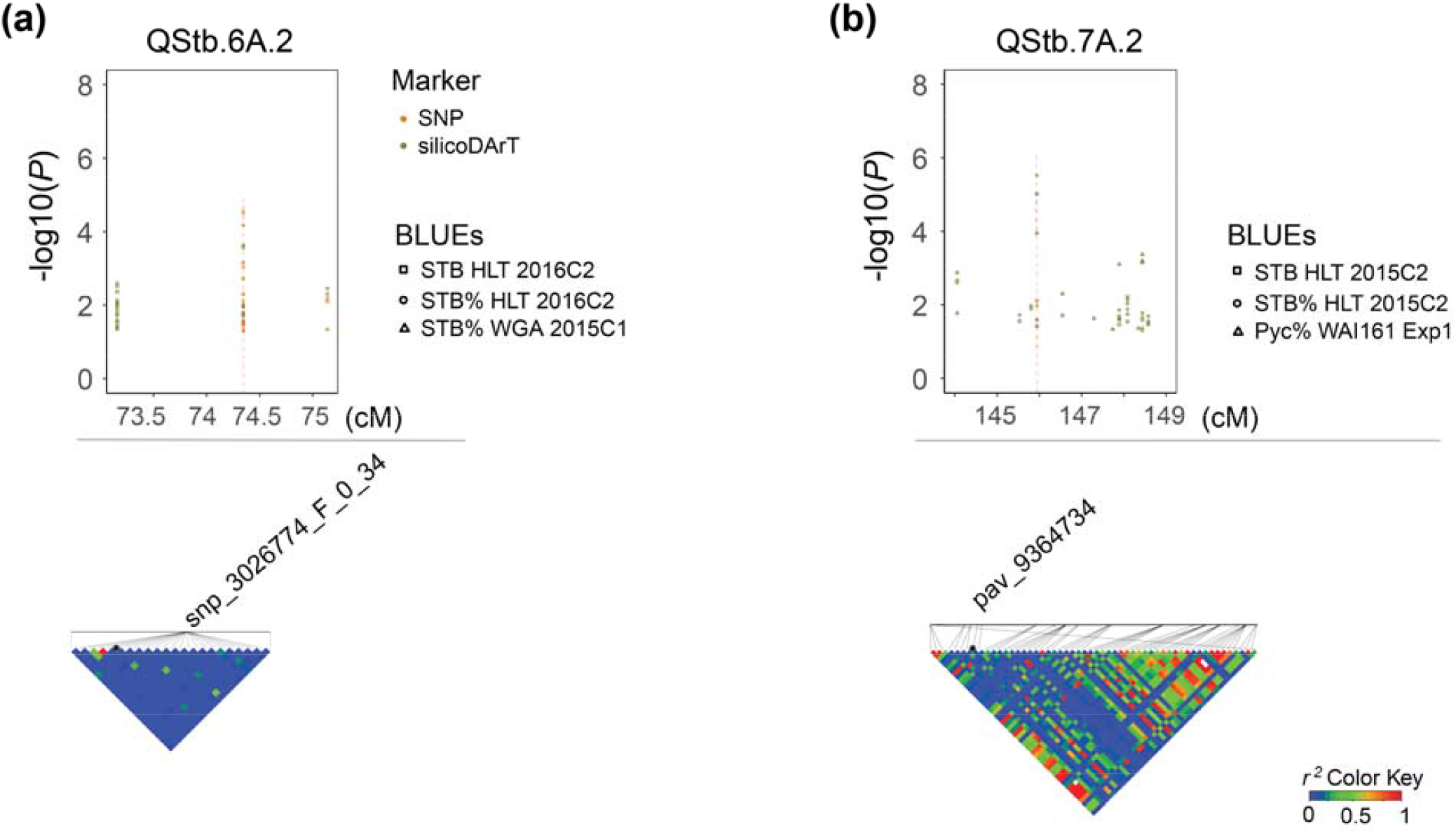
Manhattan plots and corresponding linkage disequilibrium *r*^*2*^ patterns for QStb.wai.6A.2 (**a**) associated with adult-plant resistance, and QStb.wai.7A.2 (**b**) associated with the multiple stage resistance. The upper part of the graph shows -log(*P*) value plots of marker-trait associations with detected BLUEs (FDR < 0.3). Representative SNP and silicoDArT markers and corresponding local LD *r*^*2*^ value patterns are presented in the lower part of the graph. Blue color indicates low linkage disequilibrium while red color indicates high linkage disequilibrium.

**Figure 5.**
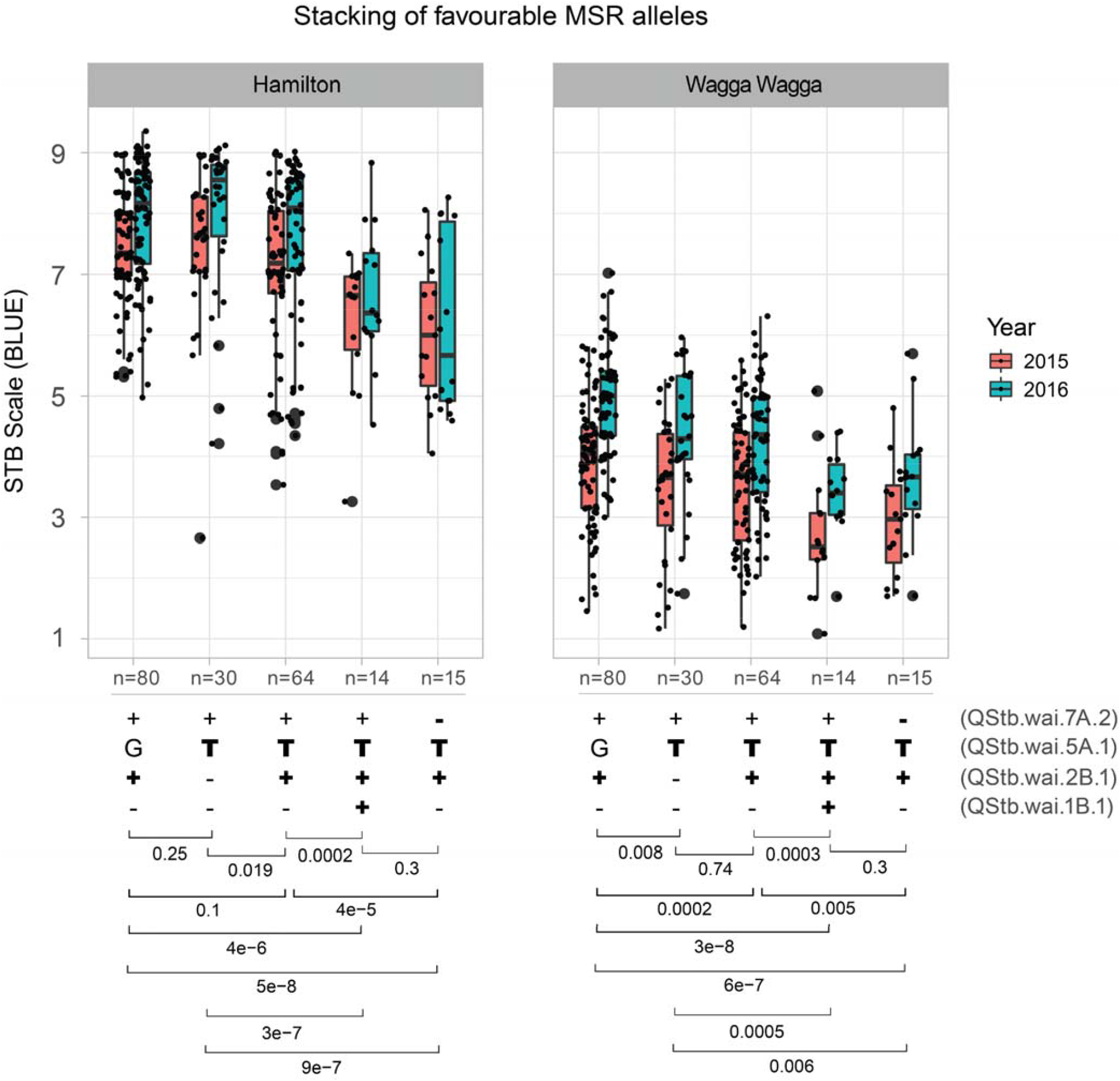
Box plot analysis of four QTLs associated with multiple-stage resistance (MSR) using their representative SNP/silicoDArT markers. Four types of stacking of alleles that existed in the AusSTB panel were shown in the lower part of the figure. Favorable alleles of QTLs are highlighted in bold. Significant *p*-values were shown at the bottom, generated by multiple-group Wilcoxon test.

### Candidate genes in the QTL regions

Physical genomic regions of thirteen QTLs were BLAST against 341 cloned genes, but only 10 of them were found to have candidate R genes ranging from 1 to 36 (Table S4). NBS (Nucleotide-site Binding) like R genes were the most abundant in seven of the ten QTLs, with the number varying from 2 to 23. Only one TaWAKL (Wall-Associated Kinase-like) R gene was found present in the QTL of QStb.wai.1A.1 associated with the WAI251-specific resistance, while only one RLK (Plant Receptor Kinase) like R gene was present in the region of QStb.wai.6A.2 responsible for APR (Table 2).

## Discussion

### Influence of marker type on the GWAS analysis

The silicoDArT markers performed slightly superior to SNP markers in detecting QTLs. Two to five times more abundance of silicoDArTs than SNPs (Table 1) increased the coverage of makers on the genome (Table 1), possibly explaining why five QTLs were detected by silicoDArTs in comparison to three QTLs by SNPs. In addition, the differences between LD distance decay for the silicoDArT and SNP reported in this study is comparable to previous studies (Cavanagh et al. 2013; Chao et al. 2010; Ovenden et al. 2017; Wang et al. 2014). The more rapid LD decay in the silicoDArTs may have helped to increase the detection of QTLs in smaller regions, therefore increasing the resolution. However, no major difference between silicoDArTs and SNPs was evident in the analysis of population structure. This is possibly due to the existence of large blocks of LD in the AusSTB panel. The recent completion of 1000 exome sequencing of wheat provides another way to enrich LD blocks using low-resolution genotyping services (He et al. 2019), which potentially increases the power to detect QTLs (Jordan et al. 2015; Nyine et al. 2019).

### Association between population genetic structure and STB resistance

The genetic characterization of the 273 bread wheat accessions divided the AusSTB panel into seven subpopulations with closely genomic related accessions. The results from the STRUCTURE analysis revealed different levels of admixtures across different subpopulations (Fig. 1b and 1c), reflecting the frequent germplasm exchanges over many years among wheat breeding programs from NSW, VIC, SA, and WA in Australia. However, high levels of resistance were observed to have a high correlation (*r* > 0.3) with slow-maturing accessions (low Zadoks scale, Table S2) at the adult-plant stage. A high correlation between these characteristics was also observed in several other genetic studies of *Z. tritici* resistance (Arraiano and Brown 2006; Dreisigacker et al. 2015; Gerard et al. 2017; Kidane et al. 2017; Muqaddasi et al. 2019; Naz et al. 2015). These results imply that winter-type or slow-growing accessions are inclined to having better *Z. tritici* resistance than the spring type or fast-growing accessions in the Australian environment. This could be due to the importance of STB resistance in the higher rainfall target environments that these longer season wheat cultivars are developed for, so breeding strategies for these types of cultivars favour the accumulation of STB resistance alleles. This correlation could also indicate that growth stage-dependent resistant QTLs are important in wheat plants at the tillering stage and booting stage, however, few studies have conducted such an exploration.

### Known resistance QTLs effectiveness revealed in Australian environments

Until now breeders targeting *Z. tritici* resistance in Australia have had limited knowledge about which resistance loci are effective in the Australian wheat gene pool, i.e. Stb2 mapped from ‘Veranopolis’ (Liu et al. 2013), Stb3 from ‘Israel 493’ (Goodwin et al. 2015), StbWW from ‘WW2449’ (Raman et al. 2009), and Stb19 from ‘Lorikeet’ (Yang et al. 2018). This GWAS study has revealed seven of the thirteen resistant QTL identified in the AusSTB panel were found to co-locate in regions previously described from international studies as containing major genes for resistance to *Z. tritici* (Brading et al. 2002; Chartrain et al. 2005a; Chartrain et al. 2005b; Chartrain et al. 2009; Cowling 2006; Cuthbert 2011; Liu et al. 2013; Raman et al. 2009). Another four QTLs identified in this study co-located at the same physical chromosome position as previously reported QTLs or within the confidence intervals of the reported QTLs (Arraiano et al. 2007; Dreisigacker et al. 2015; Goudemand et al. 2013). Some of these older reported QTLs have large regions of the chromosome associated with resistance and it is not possible to resolve if the QTLs in this study are the same as the older QTLs or novel resistance loci.

The frequencies of R alleles varied substantially in the whole population (Table 2) and subpopulations (data not shown). Five of the seedling QTLs co-located with known major R genes, which is not surprising given the use of cultivars with these R genes as parents in Australian breeding over the past 50 years. Several of the favourable allele frequencies are being maintained at high levels, such as QStb.wai.4A.1 (0.77) and QStb.wai.6A.1 (0.83), QStb.wai.7A.1 (0.88) and QStb.wai.1A.1 (0.9), which is notable considering the AusSTB panel is comprised of a wide sample of international, historic, and recent Australian cultivars and that few breeding programs have historically actively selected for seedling resistance to *Z. tritici*. These QTLs were not detected in the analysis of the adult-plant disease phenotypes. However, they must be contributing to the improved seedling-stage performance in the field to be present in such a high number of accessions in the AusSTB panel. The representative markers described here with the QTLs, enable the selection of multiple favourable alleles and enable the removal of unfavourable alleles from breeding programs (Table 2).

Two of the five QTLs identified in the adult-plant growth stages may co-locate with known major gene resistance loci. The locus QStb.wai.2B.1 is located physically close to the reported location for Stb9 (Chartrain et al. 2009) and QStb.wai.5A.1 appears to be close to the physical location reported for Stb17 (Fig. 3). The previous report of INT 6 (Yates et al. 2019) and QStb.sn.2B (Aouini 2018) also highlighted the importance of loci on chromosome 2BL for resistance. Further, the QTLs, QTL-2BL and Qstb2B_1 were also mapped from the durum wheat ‘Agili 39’and are reported to be responsible for *Z. tritici* resistance at both seedling stage and adult-plant stage using multiple isolates. It has also been suggested that this locus could be the major gene Stb9 (Aouini 2018; Ferjaoui et al. 2021).

The major gene Stb17 was sourced from synthetic bread wheat accession ‘SH M3’, and reportedly accounts for 12-32% of the adult-plant resistance (Tabib Ghaffary et al. 2012). When the available sequences from the report of Tabib Ghaffary et al. (2012) are BLAST searched against the IWGSC reference genome, Stb17 is possibly located in the region of 520-560 Mb (data not shown), while QStb.wai.5A.1 was in the region of 570-590 Mb (Table S3), close to where the *Vrn-A1* is located (IWGSC et al. 2018). In addition, QStb.wai.5A.1 with a resistant frequency of 0.56, was observed to be highly associated with low-Zadoks, APR, and WAI251-specific phenotypes, and appears to be co-located with the previously reported loci QStb.cim-5AL-2 (Dreisigacker et al. 2015) and QStb.B22-5A.a (Naz et al. 2015). It is possible that these reported loci are the gene *Vrn-A1a*, as 162 out of 273 (0.59) accessions in the AusSTB panel were identified as having *Vrn-A1a*. The *Vrn-A1a* gene, which encodes a MADS-box transcription factor 14-like protein (Yan et al. 2003), may have pleiotropic effects on the plant growth and the plant defence on different plant pathogens including *Z. tritici*, Fusarium Head Blight (Xu et al. 2019), tan spot and Septoria nodorum (Hu et al. 2019). However, the association of this locus with a seedling resistance phenotype to the WAI251 isolate suggests otherwise, these loci may be a new gene very close to *Vrn-A1a*. Some probable candidate genes at this locus include a plant receptor kinase or WAKL gene (Table S4).

### QTLs associated with APR

Generally, APR is considered preferable in breeding programs because of the flexible use in the IDM systems (Wellings et al. 2012). The putative new locus QStb.wai.6A.2 was detected across multiple sites and years, and the probable physical location for this QTL spanned from 440 Mb to 615 Mb on the Chinese Spring reference genome (S Table 3). Above the region of QStb.wai.6A.2, QTLs INT 10 and INT 11 were detected based on single year data and was positioned at 411-425 Mb. These QTL are reported to account for the control of *Z. tritici* pycnidia density within lesions at the adult-plant stage, and they were also thought to co-locate with the major gene Stb15 (Yates et al. 2019). Another QTL, QStb.teagasc-6A.2 (534-580 Mb) was associated with the resistance of flag-1 leaves to *Z. tritici* from a single-year field phenotype obtained from an Multi-parent Advanced Generation Inter-Cross (MAGIC) population (Riaz et al. 2020). It is possible that the locus QStb.teagasc-6A.2 may be the same as the QStb.wai.6A.2 as the probable physical locations of these loci overlap by approximately 50 Mb on chromosome 6A. However, accessions that carry the resistant alleles from both this study and Riaz et al. (2020) would need to be compared to determine if this is the case.

The genetic control of APR is provided by QTLs that are most effective between tillering and full head emergence, and not necessarily at the seedling stage (Ellis et al. 2014; Wellings et al. 2012). Two potential issues here might impede the utilization of APR-QTLs as breeding targets for resistance. Firstly, if a cultivar is only relying on the combination of several APR-QTLs, it is likely to be vulnerable to disease infection at the seedling stage. In cooler climate and higher rainfall areas of the south-eastern Australian, the *Zymoseptoria* population can start releasing ascospores and infecting seedlings sown in the early planting window from February to May. Secondly, if a cultivar relies only on a single APR gene, the *Z. tritici* population infecting a crop of that cultivar will only need to mutate once to overcome the resistance. One solution to overcome these two issues is to stack a combination of 2-4 major genes resistant at the seedling stage and at the adult-plant stage together in a cultivar. Before this can be attempted, the limited resource of *Z. tritici* resistance will need to be expanded. Only 24 major genes for *Z. tritici* resistance have been reported to date (Aouini 2018; Langlands-Perry et al. 2022; Tabib Ghaffary et al. 2018; Yang et al. 2018), compared to over 200 rust resistance genes (Zhang et al. 2020). The current stocks of major gene resistance are also being depleted as Australian *Z. tritici* populations evolve to overcome the effectiveness of these loci completely or partially. The loci that are known to have been overcome and are no longer effective in the Australian environment including Stb2/11/WW from ‘Veranopolis’ and ‘WW2449’, Stb3 from ‘Israel 493’, Stb4 from ‘Tadorna’, Stb6 from ‘Heraward’, Stb7 (Stb12) from ‘Currawong’, Stb14 from ‘M1696’, Stb18 from ‘Balance’ (pers. comm A. Milgate). Another possible solution to more sustainable disease resistance would be to stack major genes and APR-QTLs together. Multiple evidence suggests that combinations of different types of partial or quantitative resistance will prolong the life of *Z. tritici* resistance in cultivars, compared to a single gene of resistance (Brown 2015; Niks et al. 2015; Rimbaud et al. 2021; St Clair 2010). In this scenario, stacking of two major genes at seedling stages and two APR-QTLs into a targeted elite cultivar is not trivial, because the successful rate to capture one combination of four QTLs into a single genotype is 1/256. In comparison, the combination of two or three MSR-QTLs (1/16 or 1/64) should achieve the same level of resistance but with less breeding effort.

### QTLs associated with multi-stage resistance

The results of this study highlight the presence of QTL that provide resistance to the development of disease at multiple growth stages in plants. Multi-stage resistances can be considered different and distinct to APR, as the resistance is continuously expressed through progressive crop development stages from seedlings to grain-filling. From the point of view of resistance breeding, APR loci are attractive breeding targets for incorporation into new cultivars as they provide benefits at the flowering and grain filling stages of crop development where preservation of green leaf area has a relatively larger contribution to the final grain yield. However, before these growth stages, local transmission of *Z. tritici* inoculum is driven primarily via splash-borne pycnidiospores dispersing vertically upwards through the plant canopy from the lower layer of leaves (Eyal 1987; Robert et al. 2018). The control of *Z. tritici* from the seedling stage to booting stage, to some extent help plants reduce the amount in-crop of inoculum, which in turn alleviates the level of disease infection at later stages of plant development. MSR loci that can provide resistance (i.e., seedling-stage resistance) that reduces early levels of infection, as well as APR-type resistance that protects green leaf area at later growth stages, should be attractive breeding targets for cultivar development, particularly when they can be used in conjunction with other resistance loci for either seedling or APR.

This study introduces the concept of multi-stage resistance as a distinctive classification of loci that confer disease resistance at both seedling and some of the adult growth stages. So far since the first report of the major gene Stb5 in 2001(Arraiano et al. 2001), forty-nine genetic studies have reported over 300 genes/QTLs associated with *Z. tritici* resistance, including approx. 200 APR-QTLs and 76 seedling-stage/isolate-(non)specific genes/QTLs (Table S6). Among those, approx. twenty previously reported QTLs may fall into the MSR-QTL class of resistance (Table S5). For instance, Stb1, Stb4, Stb5, Stb6, Stb16q, and Stb18 might be the major genes known to provide MSR (Brown et al. 2015). Additionally, 16 reported QTLs with minor effects might also provide MSR (Table S5), including those discovered from five segregating populations (Aouini 2018; Eriksen et al. 2003; Piaskowska et al. 2021; Tabib Ghaffary et al. 2012; Tabib Ghaffary et al. 2011) and two association mapping populations (Goudemand et al. 2013; Louriki et al. 2021). In this study, the four identified MSR-QTLs are likely to provide resources for the development of *Z. tritici* resistant cultivars both in Australia and globally. In comparison to previously published studies, our detected MSR-QTL QStb.wai.1B.1 was co-located with QStb.cim-1BS (Dreisigacker et al. 2015), and the locus of QStb.wai.2B.1 co-located with QTL-2BL and Qstb2B_1 (Aouini 2018; Ferjaoui et al. 2021) and the major gene Stb9 (Chartrain et al. 2009). While the locus of QStb.wai.5A.1 co-located with QStb.cim-5AL-2 (Dreisigacker et al. 2015) and QStb.B22-5A.a (Naz et al. 2015). Finally, QStb.wai.7A.2 is a putative new MSR-QTL located in the distal chromosome of 7A, roughly located in 705-720 Mb based on the CS physical map (Table S4). This locus is close to but not overlapping the APR-QTLs, QStb.NS-7A (Vagndorf et al. 2017) and MQTL24 (Goudemand et al. 2013), which are estimated to be located between 680-700 Mb on the CS physical map.

The MSR-QTLs reported in this study were shown to significantly reduced disease levels when at least three were in combination. The MSR-QTL will provide a new resource for *Z. tritici* resistance breeding, although further work will be required to ascertain the genetic architecture of the QTL and validate them across multiple genetic backgrounds. These QTL are likely to be high quality targets for the development of molecular markers and target genome sequencing to identify and clone the underlying resistant genes.

## Conclusions

In summary, the study discovered eight QTLs responsible for the seedling resistance, one putative new QTL QStb.6A.2 responsible at the adult-plant stage, four QTLs responsible for MSR including the putative new QStb.wai.7A.2 at multiple stages. The underlying function and how they are acting on the pathogen during infection warrant further detailed studies as they may hold the key to more durable quantitative resistance gene of combinations.

## Supplementary Information

The online version contains supplementary material available at.

## Author contribution statement

AM conceived of the study and coordinated its design and execution. NY and AM conducted the experiments and collected the data from the experiments. NY analysed the data and wrote the draft. NY, BO, BB, MM, PS, and AM reviewed and wrote the manuscript. All authors read and approved the final manuscript.

## Acknowledgments

The Authors thank M McCaig, T Goldthorpe and M Spackman for their expert technical contributions. The authors also thank Dr A Kilian from DArT Pty Ltd for detailed advice on genotyping and GWAS analysis. This study was conducted as a co-investment between the NSW Department of Primary Industries, the Australian National University and the Grain Research and Development Corporation, under project DAN00203 of the Grains, Agronomy and Pathology Partnership.

## Data availability

All data, models, or code generated or used during the study are available from the corresponding author by request.

## Compliance with ethical standards

Conflict of Interest The authors declare that the research was conducted in the absence of any commercial or financial relationships that could be construed as a potential conflict of interest.

## References

Adhikari TB, Cavaletto JR, Dubcovsky J, Gieco JO, Schlatter AR, Goodwin SB (2004) Molecular Mapping of the Stb4 Gene for Resistance to Septoria tritici blotch in Wheat. Phytopathology 94:1198–1206

Alemu A, Brazauskas G, Gaikpa DS, Henriksson T, Islamov B, Jorgensen LN, Koppel M, Koppel R, Liatukas Z, Svensson JT, Chawade A (2021) Genome-wide association analysis and genomic prediction for adult-plant resistance to Septoria tritici blotch and powdery mildew in winter wheat. Front Genet 12:661742

Aouini L (2018) Durum wheat and Septoria tritici blotch: genes and prospects for breeding. Wageningen University

Arraiano L, Balaam N, Fenwick P, Chapman C, Feuerhelm D, Howell P, Smith S, Widdowson J, Brown J (2009) Contributions of disease resistance and escape to the control of Septoria tritici blotch of wheat. Plant Pathol 58:910–922

Arraiano L, Brown J (2006) Identification of isolate□ specific and partial resistance to Septoria tritici blotch in 238 European wheat cultivars and breeding lines. Plant Pathol 55:726–738

Arraiano L, Chartrain L, Bossolini E, Slatter H, Keller B, Brown J (2007) A gene in European wheat cultivars for resistance to an African isolate of Mycosphaerella graminicola. Plant Pathol 56:73–78

Arraiano L, Worland A, Ellerbrook C, Brown J (2001) Chromosomal location of a gene for resistance to Septoria tritici blotch (Mycosphaerella graminicola) in the hexaploid wheat’Synthetic 6x’. Theor Appl Genet 103:758–764

Arraiano LS, Brown JK (2017) Sources of resistance and susceptibility to Septoria tritici blotch of wheat. Mol Plant Pathol 18:276–292

Blake JJ, Gosling P, Fraaije BA, Burnett FJ, Knight SM, Kildea S, Paveley ND (2018) Changes in field dose–response curves for demethylation inhibitor (DMI) and quinone outside inhibitor (QoI) fungicides against Zymoseptoria tritici, related to laboratory sensitivity phenotyping and genotyping assays. Pest Manage Sci 74:302–313

Brading PA, Verstappen EC, Kema GH, Brown JK (2002) A gene-for-gene relationship between wheat and Mycosphaerella graminicola, the Septoria tritici blotch pathogen. Phytopathology 92:439–445

Brown JK (2015) Durable resistance of crops to disease: a Darwinian perspective. Annu Rev Phytopathol 53:513–539

Brown JK, Chartrain L, Lasserre-Zuber P, Saintenac C (2015) Genetics of resistance to Zymoseptoria tritici and applications to wheat breeding. Fungal Genet Biol 79:33–41

Butler D, Cullis BR, Gilmour A, Gogel B, Thompson R (2018) ASReml-R Reference Manual Version 4. University of Wollongong

Cavanagh CR, Chao S, Wang S, Huang BE, Stephen S, Kiani S, Forrest K, Saintenac C, Brown-Guedira GL, Akhunova A, See D, Bai G, Pumphrey M, Tomar L, Wong D, Kong S, Reynolds M, da Silva ML, Bockelman H, Talbert L, Anderson JA, Dreisigacker S, Baenziger S, Carter A, Korzun V, Morrell PL, Dubcovsky J, Morell MK, Sorrells ME, Hayden MJ, Akhunov E (2013) Genome-wide comparative diversity uncovers multiple targets of selection for improvement in hexaploid wheat landraces and cultivars. Proc Natl Acad Sci U S A 110:8057–8062

Chao S, Dubcovsky J, Dvorak J, Luo M-C, Baenziger SP, Matnyazov R, Clark DR, Talbert LE, Anderson JA, Dreisigacker S (2010) Population-and genome-specific patterns of linkage disequilibrium and SNP variation in spring and winter wheat (Triticum aestivum L.). BMC Genomics 11:727

Chartrain L, Berry ST, Brown JKM (2005a) Resistance of wheat line Kavkaz-K4500 L.6.A.4 to Septoria tritici blotch controlled by isolate-specific resistance genes. Phytopathology 95:664–671

Chartrain L, Joaquim P, Berry S, Arraiano L, Azanza F, Brown J (2005b) Genetics of resistance to Septoria tritici blotch in the Portuguese wheat breeding line TE 9111. Theor Appl Genet 110:1138–1144

Chartrain L, Sourdille P, Bernard M, Brown JKM (2009) Identification and location of Stb9, a gene for resistance to Septoria tritici blotch in wheat cultivars Courtot and Tonic. Plant Pathol 58:547–555

Cleveland WS, Devlin SJ (1988) Locally weighted regression: an approach to regression analysis by local fitting. Journal of the American statistical association 83:596–610

Coombes N (2002) Excavating for designs: SpaDes to DiGGer, spatial design search. Australasian Genstat Conference 2002

Courtois B, Audebert A, Dardou A, Roques S, Ghneim-Herrera T, Droc G, Frouin J, Rouan L, Gozé E, Kilian A (2013) Genome-wide association mapping of root traits in a japonica rice panel. Plos One 8:e78037

Cowling SG (2006) Identification and mapping of host resistance genes to Septoria tritici blotch of wheat. University of Manitoba

Cuthbert R (2011) Molecular mapping of Septoria tritici blotch resistance in hexaploid wheat (Triticum aestivum L.)

Dooley H, Shaw MW, Mehenni-Ciz J, Spink J, Kildea S (2016) Detection of Zymoseptoria tritici SDHI-insensitive field isolates carrying the SdhC-H152R and SdhD-R47W substitutions. Pest Manage Sci 72:2203–2207

Dreisigacker S, Wang X, Martinez Cisneros BA, Jing R, Singh PK (2015) Adult-plant resistance to Septoria tritici blotch in hexaploid spring wheat. Theor Appl Genet 128:2317–2329

Ellis JG, Lagudah ES, Spielmeyer W, Dodds PN (2014) The past, present and future of breeding rust resistant wheat. Front Plant Sci 5:641

Eriksen L, Borum F, Jahoor A (2003) Inheritance and localisation of resistance to Mycosphaerella graminicola causing Septoria tritici blotch and plant height in the wheat (Triticum aestivum L.) genome with DNA markers. Theor Appl Genet 107:515–527

Evanno G, Regnaut S, Goudet J (2005) Detecting the number of clusters of individuals using the software STRUCTURE: a simulation study. Mol Ecol 14:2611–2620

Eyal Z (1987) The Septoria diseases of wheat: concepts and methods of disease management. CIMMYT

Ferjaoui S, Aouini L, Slimane RB, Ammar K, Dreisigacker S, Schouten HJ, Sapkota S, Bahri BA, M’Barek SB, Visser RG (2021) Deciphering resistance to Zymoseptoria tritici in the Tunisian durum wheat landrace accession ‘Agili39’

Francis RM (2017) pophelper: an R package and web app to analyse and visualize population structure. Molecular ecology resources 17:27–32

Gerard GS, Börner A, Lohwasser U, Simón MR (2017) Genome-wide association mapping of genetic factors controlling Septoria tritici blotch resistance and their associations with plant height and heading date in wheat. Euphytica 213

Gilmour AR, Cullis BR, Verbyla AP (1997) Accounting for natural and extraneous variation in the analysis of field experiments. J Agric Biol Environ Stat 2:269–293

Goodwin SB, Cavaletto JR, Hale IL, Thompson I, Xu SX, Adhikari TB, Dubcovsky J (2015) A new map location of gene Stb3 for resistance to Septoria tritici blotch in Wheat. Crop Sci 55:35–43

Goudemand E, Laurent V, Duchalais L, Ghaffary SMT, Kema GH, Lonnet P, Margalé E, Robert O (2013) Association mapping and meta-analysis: two complementary approaches for the detection of reliable Septoria tritici blotch quantitative resistance in bread wheat (Triticum aestivum L.). Mol Breed 32:563–584

Hagerty CH, Graebner RC, Sackett KE, Mundt CC (2017) Variable competitive effects of fungicide resistance in field experiments with a plant pathogenic fungus. Ecol Appl 27:1305–1316

He F, Pasam R, Shi F, Kant S, Keeble-Gagnere G, Kay P, Forrest K, Fritz A, Hucl P, Wiebe K, Knox R, Cuthbert R, Pozniak C, Akhunova A, Morrell PL, Davies JP, Webb SR, Spangenberg G, Hayes B, Daetwyler H, Tibbits J, Hayden M, Akhunov E (2019) Exome sequencing highlights the role of wild-relative introgression in shaping the adaptive landscape of the wheat genome. Nat Genet 51:896–904

Hu W, He X, Dreisigacker S, Sansaloni CP, Juliana P, Singh PK (2019) A wheat chromosome 5AL region confers seedling resistance to both tan spot and Septoria Nodorum Blotch in two mapping populations. The Crop Journal 7:809–818

IWGSC; Appels R, Eversole K, Feuillet C, Keller B, Rogers J, Stein N, al. e (2018) Shifting the limits in wheat research and breeding using a fully annotated reference genome. Science 361

Jordan KW, Wang S, Lun Y, Gardiner LJ, MacLachlan R, Hucl P, Wiebe K, Wong D, Forrest KL, Consortium I, Sharpe AG, Sidebottom CH, Hall N, Toomajian C, Close T, Dubcovsky J, Akhunova A, Talbert L, Bansal UK, Bariana HS, Hayden MJ, Pozniak C, Jeddeloh JA, Hall A, Akhunov E (2015) A haplotype map of allohexaploid wheat reveals distinct patterns of selection on homoeologous genomes. Genome Biol 16:48

Kidane YG, Hailemariam BN, Mengistu DK, Fadda C, Pe ME, Dell’Acqua M (2017) Genome-wide association study of Septoria tritici blotch resistance in Ethiopian durum wheat landraces. Front Plant Sci 8:1586

Kourelis J, van der Hoorn RAL (2018) Defended to the Nines: 25 years of resistance gene cloning identifies nine mechanisms for R protein function. Plant Cell 30:285–299

Lagudah ES, Krattinger SG, Herrera-Foessel S, Singh RP, Huerta-Espino J, Spielmeyer W, Brown-Guedira G, Selter LL, Keller B (2009) Gene-specific markers for the wheat gene Lr34/Yr18/Pm38 which confers resistance to multiple fungal pathogens. Theor Appl Genet 119:889–898

Langlands-Perry C, Cuenin M, Bergez C, Krima SB, Gélisse S, Sourdille P, Valade R, Marcel TC (2022) Resistance of the wheat cultivar ‘Renan’ to Septoria Leaf Blotch explained by a combination of strain specific and strain non-specific QTL mapped on an ultra-dense genetic map. Genes 13:100

Li H, Vikram P, Singh RP, Kilian A, Carling J, Song J, Burgueno-Ferreira JA, Bhavani S, Huerta-Espino J, Payne T, Sehgal D, Wenzl P, Singh S (2015) A high density GBS map of bread wheat and its application for dissecting complex disease resistance traits. BMC Genomics 16:216

Liu Y, Zhang L, Thompson IA, Goodwin SB, Ohm HW (2013) Molecular mapping re-locates the Stb2 gene for resistance to Septoria tritici blotch derived from cultivar Veranopolis on wheat chromosome 1BS. Euphytica 190:145–156

Louriki S, Rehman S, El Hanafi S, Bouhouch Y, Al-Jaboobi M, Amri A, Douira A, Tadesse W (2021) Identification of resistance sources and genome-wide association mapping of Septoria tritici blotch resistance in spring bread wheat germplasm of ICARDA. Frontiers in Plant Science 12

Maccaferri M, Zhang J, Bulli P, Abate Z, Chao S, Cantu D, Bossolini E, Chen X, Pumphrey M, Dubcovsky J (2015) A genome-wide association study of resistance to stripe rust (Puccinia striiformis f. sp. tritici) in a worldwide collection of hexaploid spring wheat (Triticum aestivum L.). G3 (Bethesda) 5:449–465

Mahboubi M, Talebi R, Mehrabi R, Naji AM, Maccaferri M, Kema GH (2022) Genetic analysis of novel resistance sources and genome-wide association mapping identified novel QTLs for resistance to Zymoseptoria tritici, the causal agent of Septoria tritici blotch in wheat

McDonald BA, Mundt CC (2016) How knowledge of pathogen population biology informs management of Septoria tritici blotch. Phytopathology 106:948–955

McDonald MC, Renkin M, Spackman M, Orchard B, Croll D, Solomon PS, Milgate A (2019) Rapid parallel evolution of azole fungicide resistance in Australian populations of the wheat pathogen Zymoseptoria tritici. Appl Environ Microbiol 85:e01908–01918

Milgate A (2014) Septoria tritici blotch has increased in importance in high-rainfall areas of the southern GRDC region and careful attention to fungicide strategies is required to minimise the chances of further resistance developing. Ground Cover™ Issues. GRDC, Australia, Asutralia

Muqaddasi QH, Zhao Y, Rodemann B, Plieske J, Ganal MW, Röder MS (2019) Genome-wide association mapping and prediction of adult stage blotch infection in european winter wheat via high-density marker arrays. The Plant Genome 12

Naz AA, Klaus M, Pillen K, Léon J, Miedaner T (2015) Genetic analysis and detection of new QTL alleles for Septoria tritici blotch resistance using two advanced backcross wheat populations. Plant Breeding 134:514–519

Niks RE, Qi X, Marcel TC (2015) Quantitative resistance to biotrophic filamentous plant pathogens: concepts, misconceptions, and mechanisms. Annu Rev Phytopathol 53:445–470

Nyine M, Wang S, Kiani K, Jordan K, Liu S, Byrne P, Haley S, Baenziger S, Chao S, Bowden R, Akhunov E (2019) Genotype imputation in winter wheat using first-generation haplotype map snps improves genome-wide association mapping and genomic prediction of traits. G3: Genes|Genomes|Genetics 9:125–133

Ovenden B, Milgate A, Lisle C, Wade LJ, Rebetzke GJ, Holland JB (2017) Selection for water-soluble carbohydrate accumulation and investigation of genetic× environment interactions in an elite wheat breeding population. Theor Appl Genet 130:2445–2461

Piaskowska D, Piechota U, Radecka-Janusik M, Czembor P (2021) QTL mapping of seedling and adult plant resistance to Septoria tritici blotch in winter wheat cv. Mandub (Triticum aestivum L.). Agronomy 11

Pritchard JK, Stephens M, Donnelly P (2000) Inference of population structure using multilocus genotype data. Genetics 155:945–959

Quaedvlieg W, Kema G, Groenewald J, Verkley G, Seifbarghi S, Razavi M, Gohari AM, Mehrabi R, Crous P (2011) Zymoseptoria gen. nov.: a new genus to accommodate Septoria-like species occurring on graminicolous hosts. Persoonia: Molecular Phylogeny and Evolution of Fungi 26:57

Raman R, Milgate A, Imtiaz M, Tan M-K, Raman H, Lisle C, Coombes N, Martin P (2009) Molecular mapping and physical location of major gene conferring seedling resistance to Septoria tritici blotch in wheat. Mol Breed 24:153–164

Riaz A, KockAppelgren P, Hehir JG, Kang J, Meade F, Cockram J, Milbourne D, Spink J, Mullins E, Byrne S (2020) Genetic analysis using a multi-parent wheat population identifies novel sources of Septoria tritici blotch resistance. Genes (Basel) 11

Rimbaud L, Fabre F, Papaix J, Moury B, Lannou C, Barrett LG, Thrall PH (2021) Models of plant resistance deployment. Annu Rev Phytopathol

Robert C, Garin G, Abichou M, Houles V, Pradal C, Fournier C (2018) Plant architecture and foliar senescence impact the race between wheat growth and Zymoseptoria tritici epidemics. Ann Bot 121:975–989

Rudd JJ (2015) Previous bottlenecks and future solutions to dissecting the Zymoseptoria tritici-wheat host-pathogen interaction. Fungal Genet Biol 79:24–28

Saari E, Prescott J (1975) Scale for appraising the foliar intensity of wheat diseases. Plant Disease Reporter

Savary S, Willocquet L, Pethybridge SJ, Esker P, McRoberts N, Nelson A (2019) The global burden of pathogens and pests on major food crops. Nat Ecol Evol 3:430–439

Shin J-H, Blay S, McNeney B, Graham J (2006) LDheatmap: an R function for graphical display of pairwise linkage disequilibria between single nucleotide polymorphisms. Journal of Statistical Software 16:1–10

Simón M, Ayala F, Cordo C, Röder M, Börner A (2004) Molecular mapping of quantitative trait loci determining resistance to Septoria tritici blotch caused by Mycosphaerella graminicola in wheat. Euphytica 138:41–48

Sivasamy M, Aparna M, Kumar J, Jayaprakash P, Vikas V, John P, Nisha R, Satyaprakash Sivan K, Punniakotti E (2014) Phenotypic and molecular confirmation of durable adult plant leaf rust resistance (APR) genes Lr34+, Lr46+ and Lr67+ linked to leaf tip necrosis (LTN) in select registered Indian wheat (T. aestivum) genetic stocks. Cereal Research Communications 42:262–273

St Clair DA (2010) Quantitative disease resistance and quantitative resistance Loci in breeding. Annu Rev Phytopathol 48:247–268

Stram DO, Lee JW (1994) Variance components testing in the longitudinal mixed effects model. Biometrics:1171–1177

Tabib Ghaffary SM, Chawade A, Singh PK (2018) Practical breeding strategies to improve resistance to Septoria tritici blotch of wheat. Euphytica 214

Tabib Ghaffary SM, Faris JD, Friesen TL, Visser RG, van der Lee TA, Robert O, Kema GH (2012) New broad-spectrum resistance to Septoria tritici blotch derived from synthetic hexaploid wheat. Theor Appl Genet 124:125–142

Tabib Ghaffary SM, Robert O, Laurent V, Lonnet P, Margale E, van der Lee TA, Visser RG, Kema GH (2011) Genetic analysis of resistance to Septoria tritici blotch in the French winter wheat cultivars Balance and Apache. Theor Appl Genet 123:741–754

Tang Y, Liu X, Wang J, Li M, Wang Q, Tian F, Su Z, Pan Y, Liu D, Lipka AE (2016) GAPIT version 2: an enhanced integrated tool for genomic association and prediction. The plant genome 9:plantgenome2015.2011.0120

Tavella CM (1978) Date of heading and plant height of wheat varieties, as related to septoria leaf blotch damage. Euphytica 27:577–580

Vagndorf N, Nielsen NH, Edriss V, Andersen JR, Orabi J, Jørgensen LN, Jahoor A, Pillen K (2017) Genomewide association study reveals novel quantitative trait loci associated with resistance towards Septoria tritici blotch in North European winter wheat. Plant Breeding 136:474–482

VanRaden PM (2008) Efficient methods to compute genomic predictions. J Dairy Sci 91:4414–4423

Wang S, Wong D, Forrest K, Allen A, Chao S, Huang BE, Maccaferri M, Salvi S, Milner SG, Cattivelli L, Mastrangelo AM, Whan A, Stephen S, Barker G, Wieseke R, Plieske J, International Wheat Genome Sequencing C, Lillemo M, Mather D, Appels R, Dolferus R, Brown-Guedira G, Korol A, Akhunova AR, Feuillet C, Salse J, Morgante M, Pozniak C, Luo MC, Dvorak J, Morell M, Dubcovsky J, Ganal M, Tuberosa R, Lawley C, Mikoulitch I, Cavanagh C, Edwards KJ, Hayden M, Akhunov E (2014) Characterization of polyploid wheat genomic diversity using a high-density 90,000 single nucleotide polymorphism array. Plant Biotechnol J 12:787–796

Wellings C, Park RF, Simpfendorfer S, Wallwork H, Shankar M (2012) Adult Plant Resistance. Fact Sheet. GRDC, https://grdc.com.au/

Würschum T, Leiser WL, Longin CFH, Bürstmayr H (2017) Molecular genetic characterization and association mapping in spelt wheat. Plant Breeding 136:214–223

Xu K, He X, Dreisigacker S, He Z, Singh PK (2019) Anther Extrusion and its association with Fusarium Head Blight in CIMMYT wheat germplasm. Agronomy 10

Yang N, McDonald MC, Solomon PS, Milgate AW (2018) Genetic mapping of Stb19, a new resistance gene to Zymoseptoria tritici in wheat. Theor Appl Genet 131:2765–2773

Yates S, Mikaberidze A, Krattinger SG, Abrouk M, Hund A, Yu K, Studer B, Fouche S, Meile L, Pereira D, Karisto P, McDonald BA (2019) Precision phenotyping reveals novel loci for quantitative resistance to Septoria tritici blotch. Plant Phenomics 2019:1–11

Zadoks JC, Chang TT, Konzak CF (1974) A decimal code for the growth stages of cereals. Weed Res 14:415–421

Zhang J, Zhang P, Dodds P, Lagudah E (2020) How target-sequence enrichment and sequencing (tenseq) pipelines have catalyzed resistance gene cloning in the wheat-rust pathosystem. Front Plant Sci 11:678

Zhang Z, Ersoz E, Lai C-Q, Todhunter RJ, Tiwari HK, Gore MA, Bradbury PJ, Yu J, Arnett DK, Ordovas JM, Buckler ES (2010) Mixed linear model approach adapted for genome-wide association studies. Nature Genetics 42:355–360

Zwart RS, Thompson JP, Milgate AW, Bansal UK, Williamson PM, Raman H, Bariana HS (2010) QTL mapping of multiple foliar disease and root-lesion nematode resistances in wheat. Mol Breeding 26:107–124

