## Supplementary material for "Multi-stage resistance to *Zymoseptoria tritici* revealed by GWAS in an Australian bread wheat (*Triticum aestivum* L.) diversity panel": Supp Figures: AusSTB_ Supplementary_Figures.pptx

### Slide 1
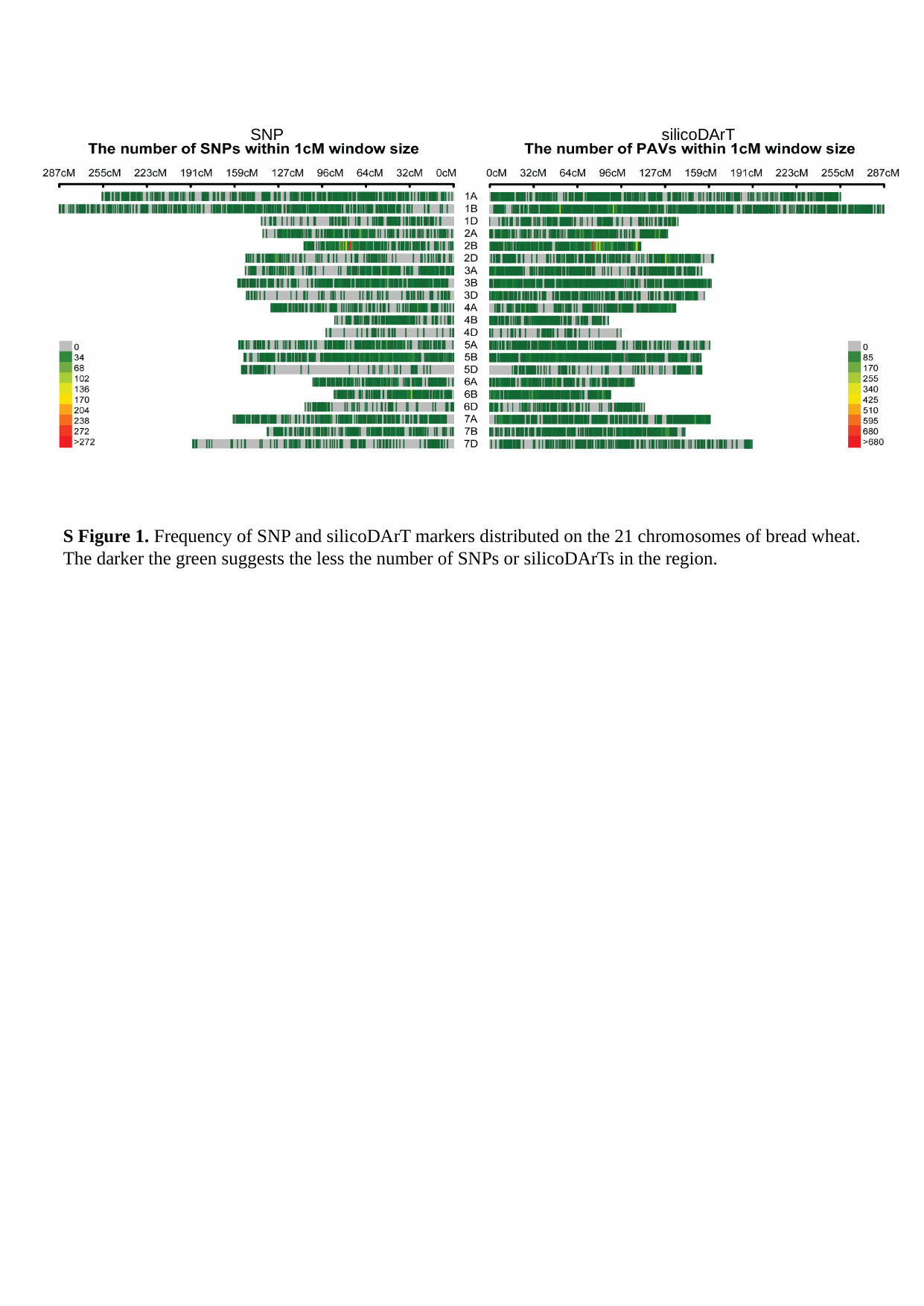

SNP
silicoDArT
S Figure 1. Frequency of SNP and silicoDArT markers distributed on the 21 chromosomes of bread wheat. The darker the green suggests the less the number of SNPs or silicoDArTs in the region.

### Slide 2
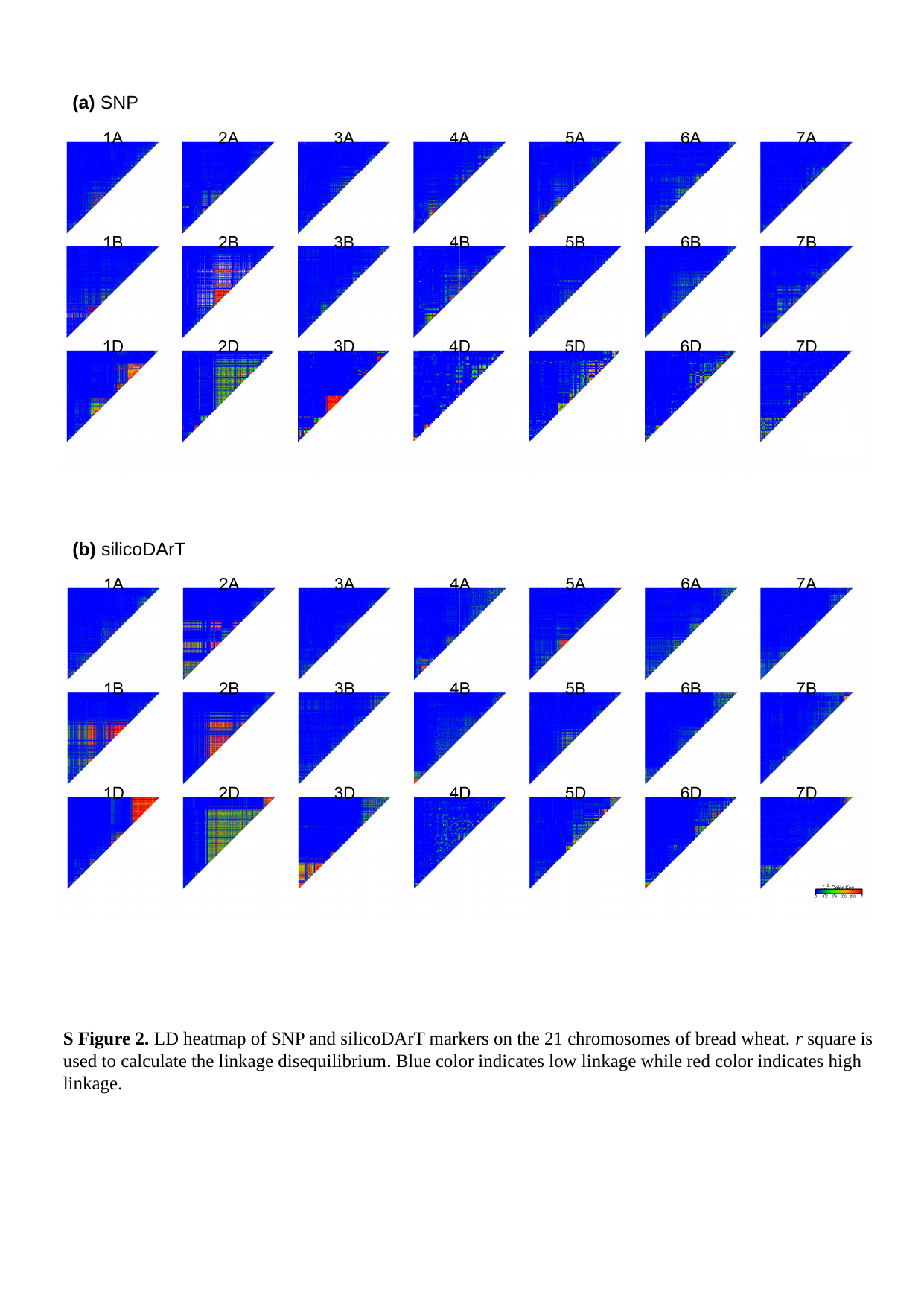

(a) SNP
(b) silicoDArT
r ^2^
S Figure 2. LD heatmap of SNP and silicoDArT markers on the 21 chromosomes of bread wheat. r square is used to calculate the linkage disequilibrium. Blue color indicates low linkage while red color indicates high linkage.

### Slide 3
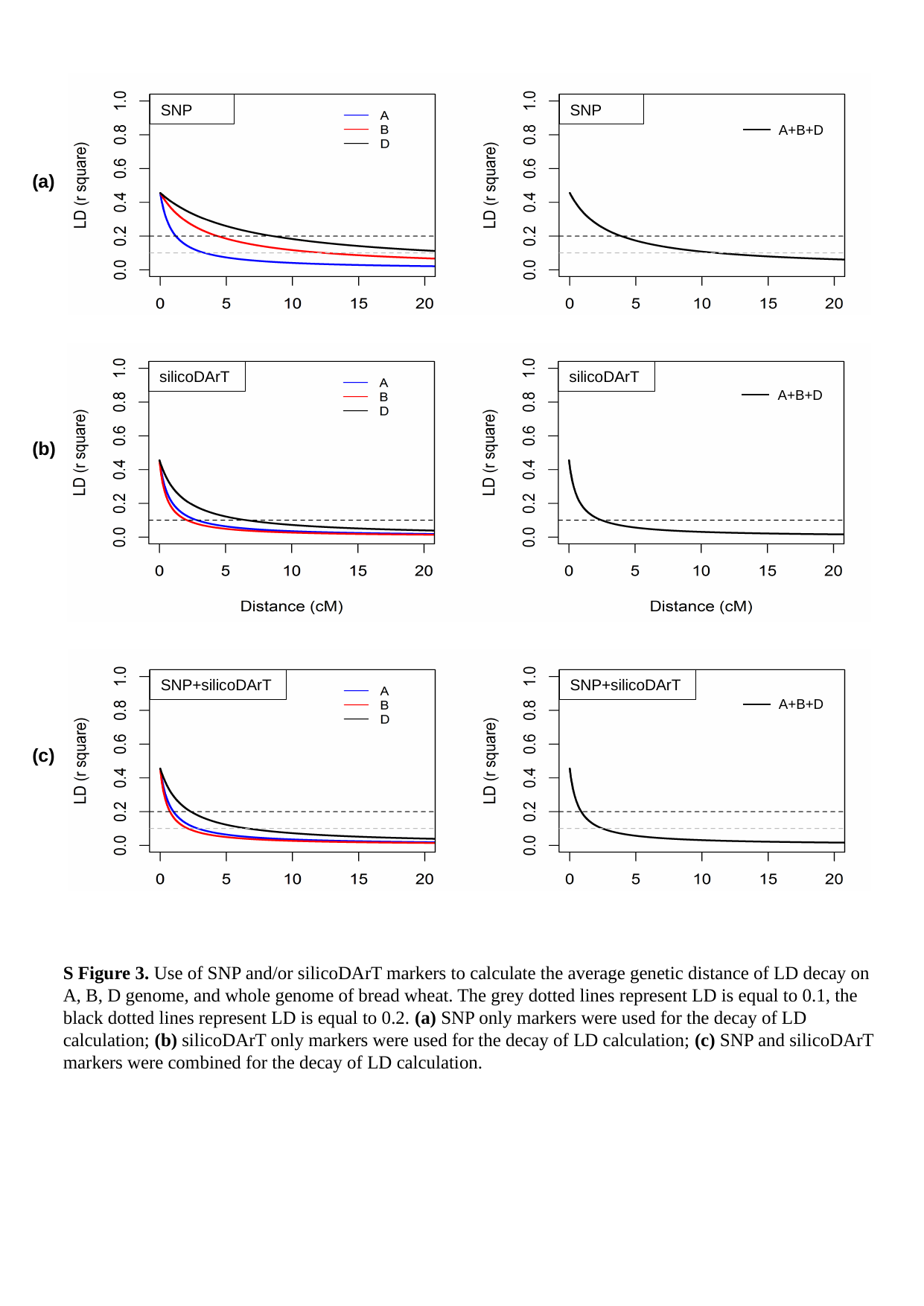

SNP
SNP
A+B+D
(a)
silicoDArT
silicoDArT
A+B+D
(b)
SNP+silicoDArT
SNP+silicoDArT
A+B+D
(c)
S Figure 3. Use of SNP and/or silicoDArT markers to calculate the average genetic distance of LD decay on A, B, D genome, and whole genome of bread wheat. The grey dotted lines represent LD is equal to 0.1, the black dotted lines represent LD is equal to 0.2. (a) SNP only markers were used for the decay of LD calculation; (b) silicoDArT only markers were used for the decay of LD calculation; (c) SNP and silicoDArT markers were combined for the decay of LD calculation.

### Slide 4
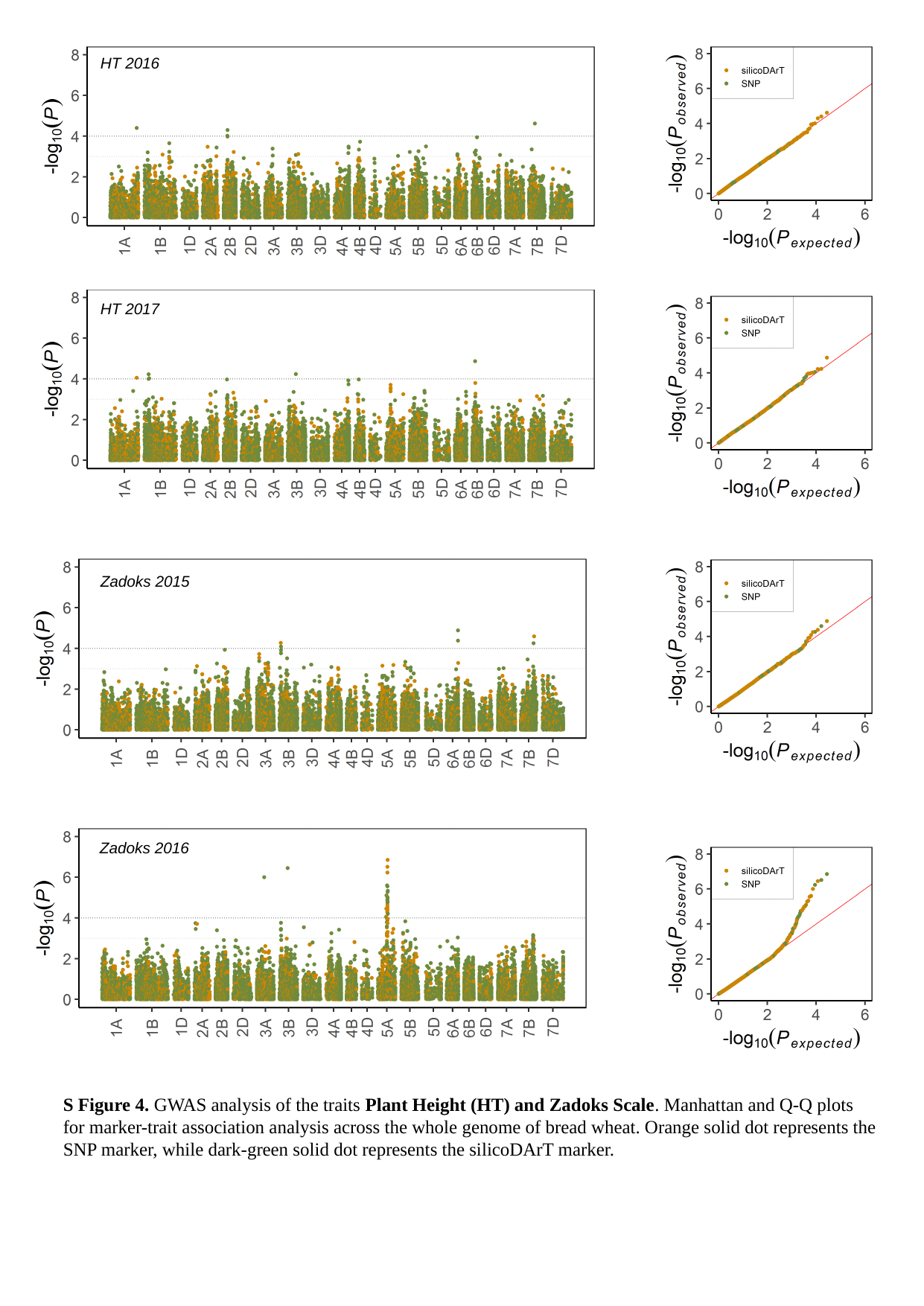

HT 2016
HT 2017
Zadoks 2015
Zadoks 2016
S Figure 4. GWAS analysis of the traits Plant Height (HT) and Zadoks Scale. Manhattan and Q-Q plots for marker-trait association analysis across the whole genome of bread wheat. Orange solid dot represents the SNP marker, while dark-green solid dot represents the silicoDArT marker.
